# Inflammation-induced mitochondrial and metabolic disturbances in sensory neurons control the switch from acute to chronic pain

**DOI:** 10.1101/2022.08.29.505682

**Authors:** Hanneke L.D.M. Willemen, Patrícia Silva Santos Ribeiro, Melissa Broeks, Nils Meijer, Sabine Versteeg, Jędrzej Małecki, Pål Ø. Falnes, Judith Jans, Niels Eijkelkamp

## Abstract

Pain often persists in patients with inflammatory diseases, even when the inflammation has subsided. The molecular mechanisms leading to this failure in resolution of inflammatory pain and the transition to chronic pain are poorly understood. Mitochondrial dysfunction in sensory neurons has been linked to chronic pain, but its role in resolution of inflammatory pain is unclear.

Transient inflammation causes neuronal plasticity, called hyperalgesic priming, which impairs resolution of hyperalgesia induced by a subsequent inflammatory stimulus. We identified that hyperalgesic priming in mice caused disturbances in mitochondrial respiration, oxidative stress, and redox balance in dorsal root ganglia (DRG) neurons. Preventing these priming-induced disturbances restored resolution of inflammatory hyperalgesia. Concurrent with these mitochondrial and metabolic changes, the expression of ATPSc-KMT, a mitochondrial methyltransferase, was increased in DRG neurons in primed mice. ATPSc-KMT overexpression in DRG neurons of naive mice induced similar mitochondrial and metabolic changes as observed after priming, leading to failure in pain resolution. Inhibition of mitochondrial respiration, knockdown of *ATPSCKMT* expression, or NAD^+^ supplementation were sufficient to restore resolution of inflammatory pain and prevent chronic pain development. Thus, inflammation-induced mitochondrial-dependent disturbances in DRG neurons promote failure in inflammatory pain resolution and drive the transition to chronic pain.

## Introduction

Chronic pain is a leading cause of years lived in disability, yet treatment options are limited and often induce severe side effects.[1] The current dogma is that pain resolution is the consequence of the dissipation of the drivers that induced the pain. However, in 12-30% of rheumatic arthritis patients pain persists while they have minimal joint inflammation or are in remission.[2] Accumulating evidence indicates that pain resolution after tissue damage or inflammation is not passive, but rather an active process that involves endogenous pain-resolution mechanisms.[3] Failed pain resolution pathways may lead to the transition from acute to chronic pain. However, the molecular mechanisms that contribute to failure in pain resolution are not well understood.

Mitochondria play a crucial role in maintaining neuronal homeostasis by ensuring metabolic functions and energy production in the form of adenosine triphosphate (ATP), via oxidative phosphorylation (OXPHOS).[4] Moreover, mitochondria are essential to regulate multiple cellular processes, such as calcium homeostasis, ion channel activity and reactive oxygen species (ROS) signaling.[4] Deficits in mitochondrial functions are linked to chronic pain. In humans, approximately 70% of the patients with heritable mitochondrial diseases have chronic pain.[5] For example, a genetic polymorphism in the mitochondrial 16S rRNA gene (gene MT-RNR2) increases the risk of developing fibromyalgia.[6] Similarly, several preclinical studies have linked mitochondrial dysfunction (e.g. reduced ATP-production) in sensory neurons to chronic pain in rodents, although predominantly in chemotherapy-induced chronic pain models.[7] In addition, the mitochondrial protein ATPSc-KMT (formerly named FAM173B), a lysine (K)-specific methyltransferase (MTase), influences OXPHOS activity by methylating Lys-43 of the ATP synthase c-subunit (ATPSc) and promotes chronic pain development.[8] Finally, OXPHOS in sensory neurons adapts during transient inflammatory pain and donation of mitochondria from macrophages to sensory neurons is needed to resolve inflammatory pain.^[3d]^ Thus, we hypothesize that adequate regulation of mitochondrial activity in sensory neurons is required for resolution of inflammatory pain. We here set out to better understand mechanistically how mitochondria in sensory neurons are involved in pain resolution or its failure, which could led to the transition from acute to chronic pain.

It is well-known that a peripheral inflammation induces long-lasting molecular changes in sensory neurons, a phenomenon called hyperalgesic priming.[9] These changes promote chronic pain development after a subsequent inflammatory insult, which normally causes only transient hyperalgesia in non-primed/naive conditions.[9] Thus, a priming model may not only help to identify mechanisms that promote chronic pain, but can also be viewed as a model that causes impairments in endogenous pain resolution mechanisms. Here, we tested whether a transient inflammation causes long-lasting disturbances in mitochondrial and metabolic activity in sensory neurons, and whether these are at the core of failure in resolution of inflammatory pain and development of chronic inflammatory pain.

## Results

### Transient inflammation causes persistent changes in mitochondrial respiration in the soma of sensory neurons

Carrageenan is a well-known priming stimulus that induces a transient inflammatory hyperalgesia and programs sensory neurons to respond differently to a subsequent inflammatory stimulus after carrageenan-induced hyperalgesia has resolved.^[9a, 10]^ Injection of carrageenan into the hind paw (intraplantar) of mice induced transient inflammatory mechanical hyperalgesia, assessed with the von Frey test. Hyperalgesia peaked at day 1 and resolved within 3-4 days (Figure 1A). At day 7, when mechanical hyperalgesia had completely resolved, mice received an intraplantar injection of PGE_2_, to unmask the primed state. PGE_2_-induced hyperalgesia persisted in primed mice (>4 days) and did not resolve within the measured time window. In contrast, in non-primed vehicle-injected mice, the PGE_2_-induced hyperalgesia resolved within one day (Figure 1A). To investigate if the presence of hyperalgesic priming at day 7 is concurrent with mitochondrial adaptations in dorsal root ganglia (DRG) that contain the soma of sensory neurons innervating the inflamed paw, we measured oxygen consumption rates (OCR) *ex vivo* as a measure of mitochondrial respiration. At the peak of inflammatory pain (day 1), basal mitochondrial respiration was reduced compared to baseline, but increased again when inflammatory hyperalgesia started resolving (at day 3; Figure 1A-C), consistent with previous findings.^[3d]^ Surprisingly, at day 7, when mechanical sensitivity had returned to baseline (Figure 1A), basal mitochondrial respiration was significantly increased compared to the respiratory activity at day 0 (Figure 1C). Dissection of basal mitochondrial respiration into three other separate key mitochondrial respiration parameters, using pharmacological mitochondrial complex specific inhibitors, showed that respiration due to proton leak, ATP synthesis-linked respiration, and maximal respiration were all increased at day 7 compared to mice without previous inflammation (Figure S1A). Mitochondrial mass or the expression of OXPHOS complexes (I-V) in lumbar DRG (Figure S1B-E) of primed mice were not significantly affected. Moreover, at day 7 after hyperalgesic priming, no signs of ongoing paw inflammation were detected, since mRNA expression of pro-inflammatory cytokines (*IL1-β, IL-6)* and macrophage marker (*F4/80*) were similar to naïve mice (Figure S1F). These data suggest that hyperalgesic priming causes increased mitochondrial respiration in DRG neurons, without clear persisting paw inflammation.

**Figure. 1:**
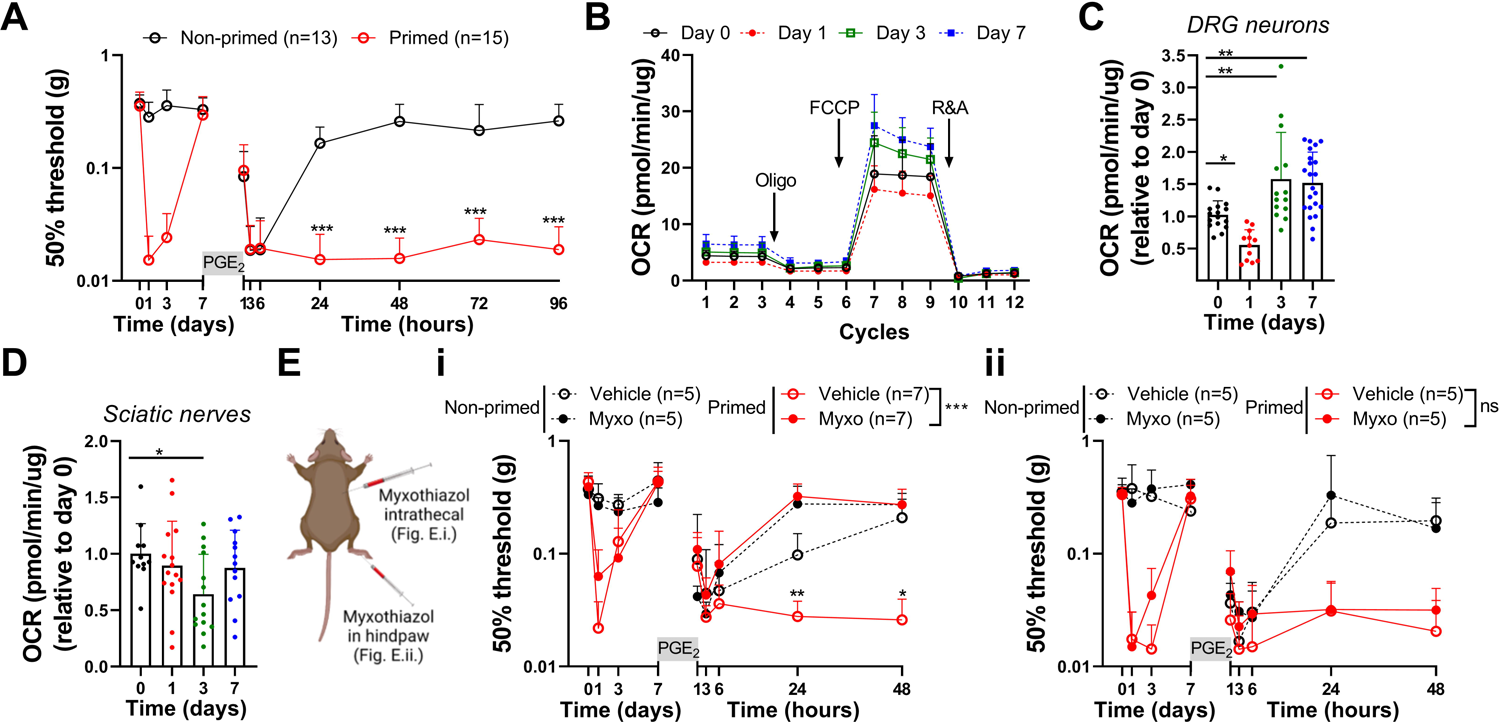
Increased mitochondrial activity in DRG neurons impairs resolution of PGE_2_-induced hyperalgesia in primed mice. **A.** Course of mechanical hyperalgesia after intraplantar injection of carrageenan (1% w/v, 5 μl, primed) or vehicle (non-primed). At day 7, mice received a subsequent intraplantar injection with PGE_2_ (100 ng/paw). **B.** OCR in DRG neuron cultures at indicated days after intraplantar carrageenan injection. OCR was measured under basal conditions and after sequential addition of oligomycin (oligo, ATP synthase inhibitor), carbonyl cyanide-p-trifluoromethoxyphenylhydrazone (FCCP, uncoupling protonophore that dissipates mitochondrial membrane potential), and a mixture of rotenone (inhibitor of Complex I) and antimycin A (inhibitor of Complex III) (R&A). **C)** Basal respiration of DRG neurons at day 0 (n=16), 1 (n=12), 3 (n=14) and 7 (n=22) after intraplantar carrageenan injection. Each dot represents the respiration measured in a well with sensory neurons. Lumbar (L3-L5) DRG from one or two mice were pooled per experiment, divided over 3–5 wells and assessed in 3 experiments. **D)** Basal respiration of sciatic nerves at day 0 (n=11), 1 (n=15), 3 (n=14) and 7 (n=13) after intraplantar carrageenan injection. Each dot is one sciatic nerve innervating an injected paw. **E)** Course of PGE_2_-induced mechanical hyperalgesia after (**i**) intrathecal or (**ii**) intraplantar injection of vehicle or myxothiazol (myxo, 50 μM) at day 7 in carrageenan-primed and non-primed mice and 15 min prior to intraplantar PGE_2_. Data are represented as mean ± SD. *P < 0.05, **P < 0.01, ***P < 0.001. Statistical analyses were performed by one-way ANOVA (C, D) followed by Dunnett’s multiple comparison test or two-way repeated measures ANOVA followed by a post-hoc Sidak’s multiple comparison test (A and E: stars indicate significance comparing primed conditions. NS = not significant).

Since energy demand may differ between the sensory neurons cell body and its axons, we also determined mitochondrial respiration in the sciatic nerves, during the course of carrageenan-induced hyperalgesia. During the peak of inflammatory hyperalgesia, basal mitochondrial respiration was similar to baseline (Figure 1D and Figure S1G). At day 3, basal mitochondrial respiration was decreased, but recovered again at day 7 to similar levels as at day 0 (Figure 1D). A similar trend was observed for proton leak, ATP synthesis-linked respiration, and maximal respiration (Figure S1H). These data indicate that after recovery from carrageenan-induced hyperalgesia, mitochondrial respiratory activity is selectively increased in the soma of sensory neurons innervating the injected paws.

### Enhanced respiratory activity in DRG neurons is linked to failure in the resolution of PGE_2_-induced hyperalgesia

We tested whether the increase in mitochondrial respiratory activity in DRG neurons contributes to the failure of resolution of PGE_2_-induced hyperalgesia in primed mice. To decrease mitochondrial respiration in lumbar DRG, mice received an intrathecal injection with myxothiazol (Complex III inhibitor of electron transport chain; ETC).[11] This administration route is effective to deliver drugs to DRG neurons[12] and significantly reduced mitochondrial respiration in DRG neurons of myxothiazol-treated mice (Figure S1I). Intrathecal administration of myxothiazol prior to PGE_2_ injection completely restored the resolution of PGE_2_-induced mechanical and thermal hyperalgesia in carrageenan-primed mice (Figure 1Ei, Figure S1J). In contrast, intraplantar injection of myxothiazol, to target nerve endings or other (local) cells, prior to PGE_2_ injection, did not restore resolution of PGE_2_-induced hyperalgesia in primed mice (Figure 1Eii). These data indicate that the enhanced mitochondrial respiratory activity in the soma of DRG neurons, but not in nerve endings, contributes to carrageenan-induced hyperalgesic priming state and associated failure in resolution of hyperalgesia, induced by a subsequent inflammatory trigger.

### Hyperalgesic priming induces disturbances in redox balance and oxidative stress in DRG

Since changes in mitochondrial respiration may affect cellular metabolism[13], we tested whether the metabolic state in lumbar DRG is affected during peripheral inflammation and after its resolution, at a time point when mice are primed. To that end, we performed a direct-infusion high-resolution mass spectrometry (DI-HRMS) on lumbar DRG lysates at various time points during carrageenan-induced hyperalgesia. With this method, we detected ∼1900 mass peaks corresponding to ∼3800 metabolites (including isomers). We performed a supervised partial least square discriminant analysis to acquire distinct metabolic profiles (Figure 2A). Subsequently, we investigated which specific metabolites are important in making those distinct clusters, followed by a metabolic pathway analysis to predict which pathways are affected by hyperalgesic priming. Explorative pathway analysis of metabolites detected in lumbar DRG at baseline (day 0) versus DRG of primed mice (day 7 after carrageenan) suggests that metabolites involved in ubiquinone synthesis, vitamin B6 metabolism, and nicotinamide metabolism are mostly affected when mice had recovered from carrageenan-induced hyperalgesia, but were primed (Figure S2A). We verified these findings by looking at the raw intensity data of metabolites involved in these pathways at day 0, 1 and 7 after carrageenan injection of the same dataset, since data-scaling and outliers may influence the outcome of the multivariate analysis. Intensities of metabolites assigned to nicotinamide metabolism, but not to ubiquinone synthesis and vitamin B6 metabolism, were significantly changed after resolution of carrageenan-induced hyperalgesia (Figure 2B, Figure S2B). These include nicotinic acid, nicotinamide riboside (NR) and quinolinic acid, which are linked to NAD^+^ biosynthesis via the Preiss-Handler pathway, salvage pathway, or the de novo synthesis of via L-tryptophan (Trp), respectively. NAD^+^ has emerged as an essential cofactor regulating mitochondrial fitness and many redox reactions.[14] Compared to day 0 (naïve mice), nicotinic acid levels were significantly increased at day 7 when mice had recovered from carrageenan-induced hyperalgesia, but not during the peak of inflammatory pain (day 1). Quinolinic acid levels were also slightly increased at day 7, but not significantly compared to DRG isolated from naive mice (p=0.07). In contrast, NR was significantly reduced at day 1 and 7 after carrageenan, compared to naive mice (Figure 2B). In an additional independent experiment, NR intensity levels were significantly reduced in DRG of primed mice that had resolved from inflammatory pain (7 days after carrageenan) compared to the peak of inflammatory pain (day 1) (Figure S2C), validating these findings. These results indicate that after resolution of a transient peripheral inflammation, when sensory neurons are primed to subsequent inflammatory triggers, formation of NAD^+^ precursors is affected in the DRG. NAD^+^ and its reduced form NADH are mainly found in three cellular pools: cytosol, nucleus and mitochondria. Cytosolic and nuclear NAD^+^ concentrations are typically similar, since NAD^+^ and NADH move freely through pores in the nuclear membrane.[15] However, mitochondrial NAD^+^ concentrations can be different from cytosolic and nuclear NAD^+^ concentrations, because the mitochondrial inner membrane is impermeable to NADH.[15] To investigate how the mitochondrial NAD^+^/NADH redox balance was affected in DRG after resolution of inflammatory pain in primed mice, we measured the 3-hydroxybutyrate/acetoacetate ratio as a proxy for the mitochondrial NAD^+^/NADH ratio.[16] At day 7 after intraplantar carrageenan administration, the 3-hydroxybutyrate/acetoacetate ratio was increased compared to day 0 (Figure 2C). These data would support an increase in the mitochondrial NAD^+^/NADH ratio, either due to higher mitochondrial NAD^+^ or lower mitochondrial NADH concentrations. The latter could be caused by reduced NADH generation in the tricarboxylic acid cycle (TCA) or increased NADH consumption by Complex I to support increased OXPHOS activity, as we had observed in primed mice (Figure 1C).

**Figure 2:**
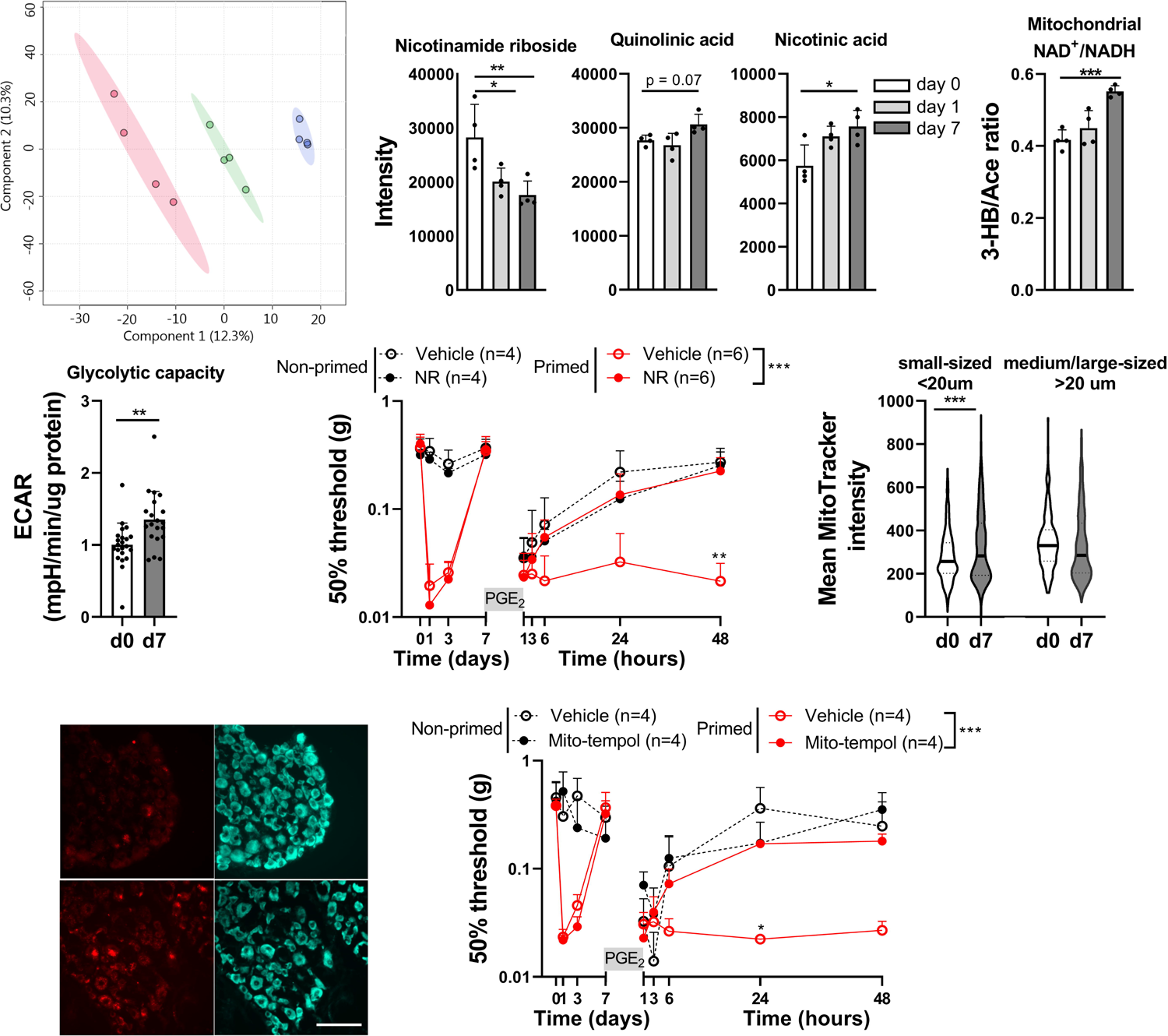
NAD^+^ supplementation and attenuation of oxidative stress restores resolution of PGE_2_-induced hyperalgesia in primed mice. **A)** Supervised partial least square discriminant analysis of DI-HRMS of lumbar DRG isolated from untreated mice (day 0) and 1 or 7 days after intraplantar carrageenan (each dot represents a metabolic signature of lumbar DRG isolated from one mouse). **B/C)** Intensity of metabolites measured with DI-HRMS **B)** involved in generation of NAD^+^ **C)** and used to measure 3-hydroxybutyrate/acetoacetate ratio as an indirect measure of mitochondrial NAD^+^/NADH ratio (n=4). **D)** Extracellular acidification rate (ECAR) in cultured DRG neurons from untreated mice (non-primed, n=22) or mice that had resolved from carrageenan-induced inflammatory hyperalgesia (primed, day 7, n=20). **E)** Course of PGE_2_-induced mechanical hyperalgesia after intraperitoneal injection with nicotinamide riboside (NR, 500 mg/kg) at day 7 in carrageenan-primed or non-primed mice and 15 min prior to intraplantar PGE_2_. **F/G)** Detection of mtROS formation in DRG neurons of untreated mice (day 0) or mice that had resolved from carrageenan-induced inflammatory hyperalgesia (day 7). mtROS was visualised by an intrathecal injection with MitoTracker Red CM-H2Xros (100 uM) that accumulates in mitochondria and generates fluorescence upon oxidation by mtROS. **F)** Mean MitoTracker Red CM-H2Xros fluorescence intensity in small and medium/large-diameter neurons at indicated days. Data visualized with a violin plot to show distribution of all data, median (black horizontal line) and quartiles (dotted line; small-diameter n= 700-1000 cells, medium/large-sized = 250-550 cells). **G)** Representative pictures of F. Neurons are visualised with neurotrace (blue; scale bar = 100 µM). **H)** Course of PGE_2_-induced mechanical hyperalgesia after intrathecal administration of mito-tempol (25 μg) at day 7 in carrageenan-primed or non-primed mice and 15 min prior to intraplantar PGE_2_. Data are represented as mean ± SD. *P < 0.05, **P < 0.01, ***P < 0.001. Statistical analyses were performed by Student’s T-test (D), one-way ANOVA (B, C, F) followed by Dunnett’s multiple comparison test or two-way repeated measures ANOVA followed by a post-hoc Sidak’s multiple comparison test (E/H, stars indicate significance comparing primed conditions).

In the cytosol, the NAD^+^/NADH redox state is strongly determined by glycolysis, which promotes the balance towards increased NADH levels.[17] The ratio between the glycolytic metabolite redox couple lactate/pyruvate, an indirect measure for cytosolic NAD^+^/NADH ratio, was unaffected in the DRG of primed mice (Figure S2D). However, the extracellular acidification rate (ECAR), a measure of anaerobic glycolysis, was significantly increased in cultured DRG neurons isolated from primed mice compared to untreated mice (Figure 2D). Overall, these data point to disturbances in glycolysis, mitochondrial respiration, nicotinamide metabolism and mitochondrial redox balance in the DRG after hyperalgesic priming.

NAD^+^ supplementation *in vivo* can mitigate enhanced glycolysis, redox disturbances and improve mitochondrial functions.[18] Moreover, systemic or oral administration of the NAD^+^ precursor NR increases NAD^+^ levels in different tissues, including nervous tissue.[19] Therefore, we tested if NR supplementation is sufficient to restore resolution of PGE_2_-induced hyperalgesia in primed mice. Indeed, NR supplementation, through an intraperitoneal injection prior to intraplantar PGE_2_-injection, prevented failure in resolution of PGE2-induced hyperalgesia in primed mice. NR supplementation did not affect PGE_2_-induced hyperalgesia in non-primed mice (Figure 2E and Figure S2E). Disturbed redox balance is often associated with oxidative stress, e.g. due to oversupply of NADH to ETC, which promotes electron leakage and mitochondrial superoxide (mtROS) production.[20] To assess mtROS production in DRG neurons, mice were injected intrathecally with MitoTrackerRedCM-H2XROS at day 7 after intraplantar carrageenan or vehicle injection.^[8a, 21]^ At day 7, when mice had recovered from inflammatory hyperalgesia, MitoTrackerRedCM-H2XROS fluorescence (Figure 2F/G) or the number of MitoTrackerRedCM-H2XROS positive small-diameter DRG neurons (<20 μm) were increased (mtROS positive/negative neurons: non-primed 35/693 (∼5%), primed 151/1021 (∼15%), p <0.0001). In medium/large-diameter neurons MitoTrackerRedCM-H2XROS fluorescence (Figure 2F/G) or the number of positive neurons (>20 μm, non-primed 25/235 (∼10%), primed 82/549 (∼15%, p = 0.0912) were not affected.[22] Pharmacological blockade of superoxide, with an intrathecal injection of the mitochondrial ROS scavenger mito-tempol[23], prior to intraplantar PGE_2_, restored resolution of PGE_2_-induced hyperalgesia in primed mice (Figure 2H, Figure S2F). In conclusion, disturbances in redox balance and oxidative stress persist in DRG neurons after resolution of inflammatory pain. Our data suggest that these disturbances lead to failing pain resolution to an inflammatory stimuli, driving the transition to chronic pain in primed mice.

### ATPSc-KMT promotes mitochondrial hyperactivity and induces hyperalgesic priming

ATPSc-KMT has recently been identified as a mitochondrial protein driving chronic inflammatory pain. ATPSc-KMT promotes mtROS formation when over-expressed and is required for efficient mitochondrial respiration.[8] Therefore, we asked whether changes in ATPSc-KMT expression may underlie the persistent mitochondrial and metabolic adaptations that cause failure of pain resolution after priming. We first evaluated *ATPSCKMT* mRNA expression in DRG during the course of carrageenan-induced hyperalgesia. *ATPSCKMT* mRNA expression was increased in lumbar DRG at day 3 and 7 after intraplantar carrageenan injection (Figure 3A). At day 7, when hyperalgesia had resolved and mice were primed, ATPSc-KMT protein level was increased in small and medium/large-diameter neurons compared to vehicle injected mice (Figure 3B/C). Thus, carrageenan increases ATPSc-KMT expression at mRNA and protein levels in DRG neurons, and these changes persists after the resolution of inflammatory hyperalgesia.

**Figure. 3:**
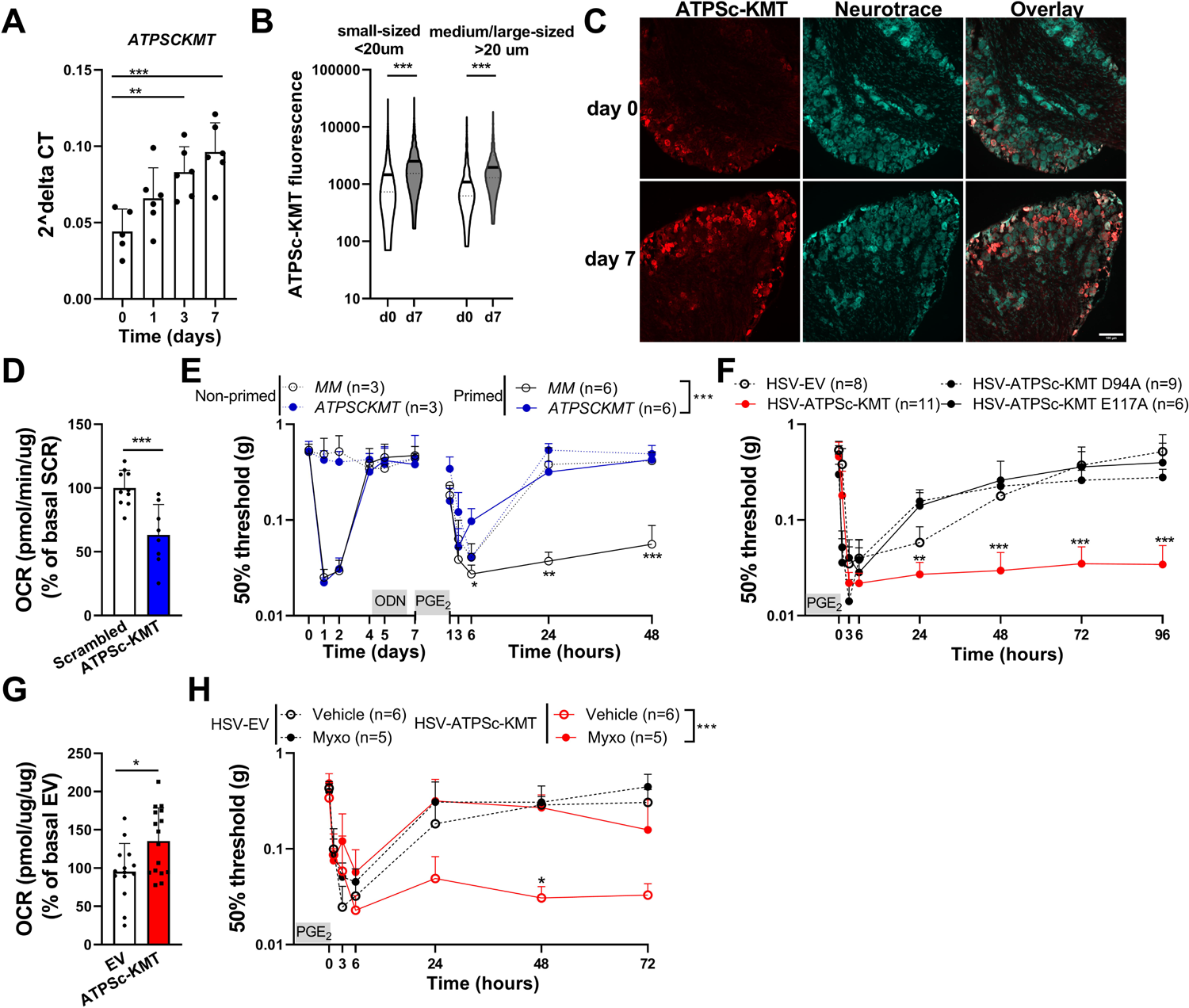
ATPSc-KMT promotes mitochondrial hyperactivity in DRG neurons and mimics hyperalgesic priming. Expression of ATPSC-KMT **A)** mRNA in the total DRG (n=6) and **B)** protein in soma of sensory neurons of lumbar (L3-L5) DRG at different days after intraplantar carrageenan. Mean ATPSC-KMT fluorescence intensity in small- and medium/large-sized neurons at indicated days. Data visualized with a violin plot to show distribution of all data with median (black horizontal line) and quartiles (dotted line; small-sized n= 1200-1300 cells, medium/large-sized = 1230-1325 cells). **C)** Example pictures of ATPSc-KMT fluorescence in DRG neurons (scale bar = 100 μM). **D)** OCR measurements in primary sensory neurons after lentiviral-mediated *ATPSCKMT* knockdown (n=8) compared to scrambled controls (n=10). **E)** Course of PGE_2_-induced mechanical hyperalgesia after intrathecal injection of *ATPSCKMT*-antisense or mismatched control (MM) oligodeoxynucleotide (ODN, 3 μg/μl, 5 μL) at day 4, 5 and 6 in carrageenan-primed and non-primed mice. **F**) Course of PGE_2_-induced mechanical hyperalgesia in mice expressing ATPSc-KMT, ATPSc-KMT enzyme-dead mutants (D94A or E117A) or EV in DRG neurons. Intraplantar HSV injections were administrated at day −3 and −1 (35.000 pfu/paw) **G)** OCR measurements after HSV-mediated ATPSc-KMT expression (n=16) in DRG neurons and HSV-empty vector (EV) expression as control (n=14). **H)** Same as F, but with intrathecal injection of myxothiazol (myxo, 50 uM) 15 min prior to intraplantar PGE_2_. Data are represented as mean ± SD. *P < 0.05, **P < 0.01, ***P < 0.001. Statistical analyses were performed by Student’s t-test (D and G), one-way ANOVA (A and B) followed by Dunnett’s multiple comparison test or two-way repeated measures ANOVA followed by a post-hoc Sidak’s multiple comparison test (E/F/H; stars indicate significance comparing carrageenan- or ATPSc-KMT-primed conditions). Primed mice by ATPSc-KMT overexpression are indicated with red bars/lines and blue bars/lines indicate *ATPSCKMT* knockdown.

Deficiency of *ATPSCKMT* impairs Complex V activity and mitochondrial respiration in HAP1 and Neuro2a (N2A) cells.^[8b]^ Here we show that *ATPSCKMT* knockdown also reduced basal mitochondrial respiration, proton leakage, ATP-driven and maximal respiration in cultured DRG neurons (Figure 3D, Figure S3A/B). To investigate whether the increased ATPSc-KMT expression in DRG neurons of primed mice prevents resolution of subsequent PGE_2_-induced hyperalgesia, we targeted ATPSc-KMT expression in the lumbar DRG by using intrathecal injections of mouse *ATPSCKMT* antisense oligodeoxynucleotide (*ATPSCKMT*-AS). To that end, mice received lumbar intrathecal injections of *ATPSCKMT*- AS at day 4, 5 and 6 after intraplantar carrageenan, a strategy which reduces *ATPSCKMT* expression and mitochondrial respiration in DRG neurons (Figure S3C).^[8a]^ *ATPSCKMT-AS* induced knockdown fully restored the resolution of PGE_2_-induced hyperalgesia in carrageenan-primed mice (Figure 3E). Resolution of PGE_2_-induced hyperalgesia in carrageenan-primed mice treated with control mismatch oligodeoxynucleotides (MM-AS) did not occur within the measured hours (Figure 3E, Figure S3D). Knockdown of *ATPSCKMT* did not affect the course of PGE_2_-induced hyperalgesia in non-primed mice (Figure 3E).

Next, we assessed whether increased ATPSc-KMT expression in DRG neurons is sufficient to mimic a hyperalgesic priming state, enhanced mitochondrial respiration and failure in resolution of PGE_2_-induced hyperalgesia. To express ATPSc-KMT *in vivo* in DRG neurons, mice were injected intraplantar with Herpes Simplex Virus (HSV) amplicons, which selectively infect primary sensory neurons.[24] Administration of HSV-ATPSc-KMT prior to intraplantar PGE_2_, prevented the resolution of PGE_2_-evoked mechanical (Figure 3F) and thermal hyperalgesia (Figure S3E). In contrast, resolution of PGE_2_-induced hyperalgesia was unaffected after expression of control empty vector (HSV-EV) or two different enzyme dead mutants, D94A (located in Motif I) and E117A (located in Post I motif)[8] (Figure 3F, Figure S3E). As anticipated, ATPSc-KMT overexpression was sufficient to increase basal respiration, proton leakage, ATP-driven, and maximal respiration in N2A cells (Figure S3F) or cultured primary sensory neurons (Figure 3G, Figure S3G/H). The increase in mitochondrial respiration was not caused by other oxygen consuming processes or cellular metabolism, as Complex II-driven respiration in isolated mitochondria was increased by ATPSc-KMT overexpression (Figure S3I). Injection of the Complex III inhibitor myxothiazol to reduce ATPSc-KMT-induced mitochondrial (hyper)activity in DRG neurons restored the resolution of PGE_2_-induced hyperalgesia in mice overexpressing ATPSc-KMT (Figure 3H, Figure S3J). These data indicate that elevated ATPSc-KMT expression in DRG neurons is sufficient to mimic the primed states of sensory neurons after a transient inflammatory stimulus, by promoting mitochondrial hyperactivity and causing failure in the resolution of PGE_2_-induced hyperalgesia.

Next, we aimed to understand how ATPSc-KMT expression increases mitochondrial activity, and impairs resolution of PGE_2_-induced inflammatory hyperalgesia. Recently, we described that Lys-43 of the ATP synthase c-subunit (Complex V of ETC), is the main substrate of ATPSc-KMT, suggesting that alterations in the methylation status of Lys-43 may regulate pain resolution.^[8b]^ However, we found that Lys-43 of the ATP synthase c-subunit was already fully methylated (trimethylated) in DRG of untreated mice (Figure S3K/L), making it unlikely that increased ATPSc-KMT activity/expression enhances mitochondrial respiratory activity through increased methylation of Lys-43 of the ATP synthase c-subunit. Indeed, when ATP synthase was fully inhibited and uncoupled from OXPHOS, *ATPSCKMT* deficient (ATPSc-KMT^-/-^) HAP1 cells still had a significantly reduced OCR compared to wild-type (WT) cells, and reduced OCR when compared to ATPSc-KMT^-/-^ cells reconstituted with WT ATPSc-KMT, but not with enzyme-dead mutant ATPSc-KMT-E117A (Figure S3M). These data suggest that ATPSc-KMT may regulate mitochondrial activity by mechanisms other than methylation of ATP synthase c-subunit.

### ATPSc-KMT expression affects cellular metabolism and disturbs redox balance

To assess whether ATPSc-KMT expression affects cellular metabolism, we performed non-quantitative DI-HRMS. An unsupervised principal component analysis (PCA) showed that ATPSc-KMT^-/-^ HAP1 cells clustered distinctly from WT cells and ATPSc-KMT^-/-^ cells reconstituted with WT ATPSc-KMT. ATPSc-KMT^-/-^ cells reconstituted with the enzyme dead mutant E117A clustered close to ATPSc-KMT deficient cells (Figure 4A). Enrichment analysis identified a group of functionally related metabolites and ranked metabolites involved in the mitochondrial ETC and the Warburg effect (high rate of glycolysis) at the top of the list (Figure S4A). Heat map analysis indicates that L-acetylcarnitine, which buffers the pool of acetyl-CoA to enter the TCA[25], was significantly reduced in ATPSc-KMT deficient cells, and normalized again by reconstitution of WT ATPSc-KMT (Figure 4B, Figure S4B). Similarly, D-glyceraldehyde 3-phosphate (GA3P) and lactic acid, the intermediate and end product metabolites of anaerobic glycolysis, respectively, were decreased two-fold in ATPSc-KMT^-/-^ cells compared to WT cells and recovered by reconstitution with WT ATPSc-KMT, but not with ATPSc-KMT-E117A (Figure 4C). The decrease of lactate was validated in a targeted quantitative analysis of TCA metabolites (Figure S4C) and in CRISPR/CAS9 generated ATPSc-KMT deficient N2A cells (Figure 4D). Decreased lactate levels are indicative of impaired glycolysis.[26] In accordance, ECAR levels, as proxy of glycolysis, were reduced in ATPSc-KMT^-/-^ cells compared to WT cells, but were restored after reconstitution with WT, but not E117A-mutated ATPSc-KMT (Figure S4D), indicating that ATPSc-KMT influences glycolysis. These data confirm our hypothesis that ATPSc-KMT expression changes cellular metabolism in HAP1 and N2A cells by promoting glycolysis and oxidative phosphorylation.

**Figure. 4:**
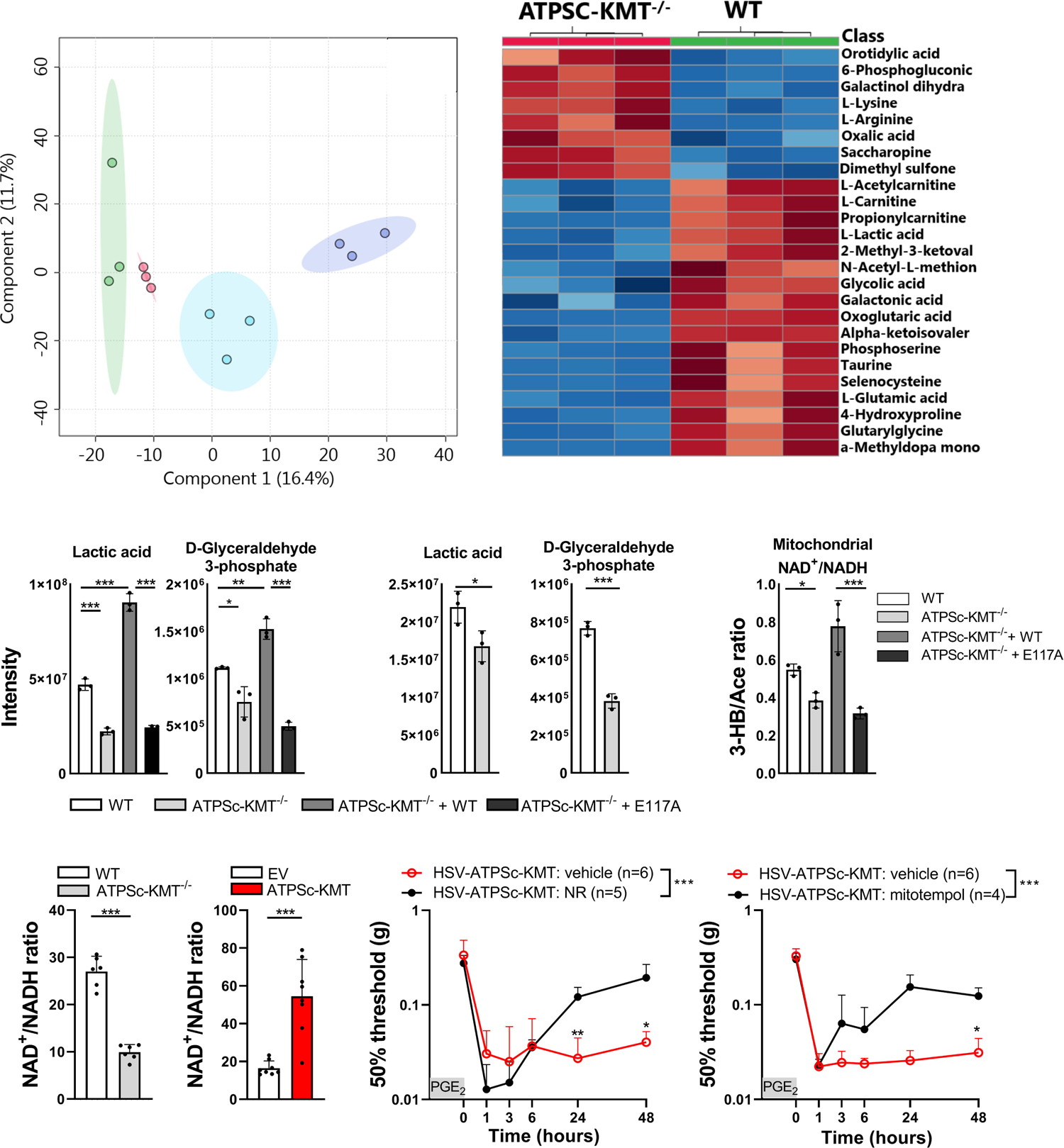
ATPSc-KMT affects cellular metabolism and redox balance. **A)** Unsupervised principle component analysis (PCA) of whole metabolomics DI-HMRS dataset of HAP1 cells expressing ATPSc-KMT (WT), cells deficient for ATPSc-KMT (KO) and KO cells reconstituted with WT (KO + WT) or with enzyme dead mutant ATPSc-KMT (KO + E117A) **B)** Heat map of metabolite levels that were significantly changed between WT and ATPSc-KMT^-/-^ cells. Blue = reduced intensity, red = increased intensity. Intensity of lactic acid and GA3P in **C)** HAP1 cells and **D)** N2A cells with and without ATPSc-KMT. **E)** Ratio of 3-hydroxybutyrate and acetoacetate which was measured with DI-HMRS and which is an indirect measure of mitochondrial NAD^+^/NADH balance. **F/G)** NAD^+^/NADH ratio calculated from direct measurement of cellular NAD^+^ and NADH in **F)** HAP1 WT and ATPSc-KMT deficient cells and **G)** N2A cells after overexpression of ATPSc-KMT or control empty vector (EV). **H)** Course of PGE_2_-induced mechanical hyperalgesia after an intraperitoneal injection of nicotinamide riboside (NR), 15 min prior to intraplantar PGE_2_ injection, in mice overexpressing ATPSc-KMT in DRG neurons. Intraplantar HSV injections were administrated at day −3 and −1 (35.000 pfu/paw) **I)** Same as H, but after intrathecal injection of mito-tempol, 15 min prior to intraplantar PGE_2_ injection. Data are represented as mean ± SD. *P < 0.05, **P < 0.01, ***P < 0.001. Statistical analyses were performed by Student’s t-test (D, F and G), one-way ANOVA (C and E) or two-way repeated measures ANOVA followed by a post-hoc Sidak’s multiple comparison test (H and I).

Since nicotinamide riboside in DRG neurons was reduced after resolution of carrageenan-induced hyperalgesia at a timepoint when ATPSc-KMT expression was increased, we investigated whether NAD^+^ precursors are affected by ATPSc-KMT. ATPSc-KMT expression and activity did not affect NR, quinolinic acid or nicotinic acid levels (Figure S4E). However, ATPSc-KMT deficiency decreased the 3-hydroxybutyrate/acetoacetate ratio, which was restored with WT ATPSc-KMT, but not with the ATPSc-KMT-E117A mutant (Figure 4E). These data suggest that ATPSc-KMT may disturb mitochondrial NAD^+^/NADH ratio. Decreased NADH levels are not favorable in mitochondria, since NADH is an important Complex I substrate for the ETC to produce ATP. In line with the indirect measurements that pointed toward a shift in the mitochondrial NAD^+^/NADH ratio (Figure 4E), direct NAD^+^ and NADH measurements with ultra-high performance liquid chromatography (UPLC) showed that the total cellular NAD^+^/NADH ratio was reduced in ATPSc-KMT deficient HAP1 cells (Figure 4F). Conversely, transient overexpression of ATPSc-KMT in N2A cells increased the cellular NAD^+^/NADH ratio, with predominately an increase in the NADH cellular pool (Figure S4F), compared to control EV-transfected cells (Figure 4G). Likely, this shift is caused by the increased consumption of NADH due to the ATPSc-KMT-induced increase in OXPHOS (Figure 3G, S3F-I). Next, we investigated whether NR supplementation was also sufficient to restore resolution of PGE_2_-induced hyperalgesia in HSV-ATPSc-KMT primed mice. An intraperitoneal injection of NR prior to intraplantar PGE_2_-injection completely prevented failure in resolution of PGE_2_-induced mechanical hyperalgesia in HSV-ATPSc-KMT primed mice (Figure 4H, S5B).

Redox balance control NADPH-requiring antioxidant pathways.[27] Thus a redox imbalance (e.g. an increased mitochondrial NAD^+^/NADH ratio) impairs the cellular anti-oxidant capacity leading to oxidative stress.^[18c, 28]^ ATPSc-KMT deficient cells had reduced mtROS levels compared to WT cells or ATPSc-KMT^-/-^ cells reconstituted with WT ATPSc-KMT, but not the enzyme dead ATPSc-KMT-E117A (Figure S5A). Intrathecal injection of the mitochondrial ROS scavenger mito-tempol, prior to intraplantar PGE_2_ injection in mice expressing HSV-ATPSc-KMT, restored the resolution of PGE_2_-induced hyperalgesia (Figure 4I, Figure S5C). These data suggest that ATPSc-KMT methyltransferase activity affects the formation of metabolites, increases oxidative stress, and disturb the redox state of sensory neurons. As a consequence, resolution of PGE_2_-induced hyperalgesia fails. Blocking the oxidative stress or NAD^+^ supplementation restores the resolution of PGE_2_-induced hyperalgesia in mice expressing HSV-ATPSc-KMT.

### NAD^+^ supplementation attenuates Complete Freund’s Adjuvant-induced chronic inflammatory pain

Since our data indicate that NR supplementation is sufficient to restore resolution of PGE_2_ hyperalgesia in carrageenan-primed mice, or mice overexpressing ATPSc-KMT, we next tested whether NR supplementation would be sufficient to resolve chronic inflammatory pain. Persistent inflammatory hyperalgesia was induced by an intraplantar injection of complete Freund’s adjuvant (CFA).^[8a, 29]^ Five days after development of CFA-induced inflammatory pain, mice received intraperitoneal (i.p.) injections with NR for 3 consecutive days. NR administration attenuated CFA-induced mechanical (Figure 5A) and thermal (Figure 5B) hyperalgesia compared to vehicle treated mice, whereas NR treatment did not have any effect on pain associated behaviors in mice that received an intraplantar vehicle injection only. Local intrathecal (i.t.) injection with NR, to target the DRG and spinal cord, but not the inflamed tissue, also resolved CFA-induced persistent inflammatory pain (Figure 5A/B).

**Figure 5:**
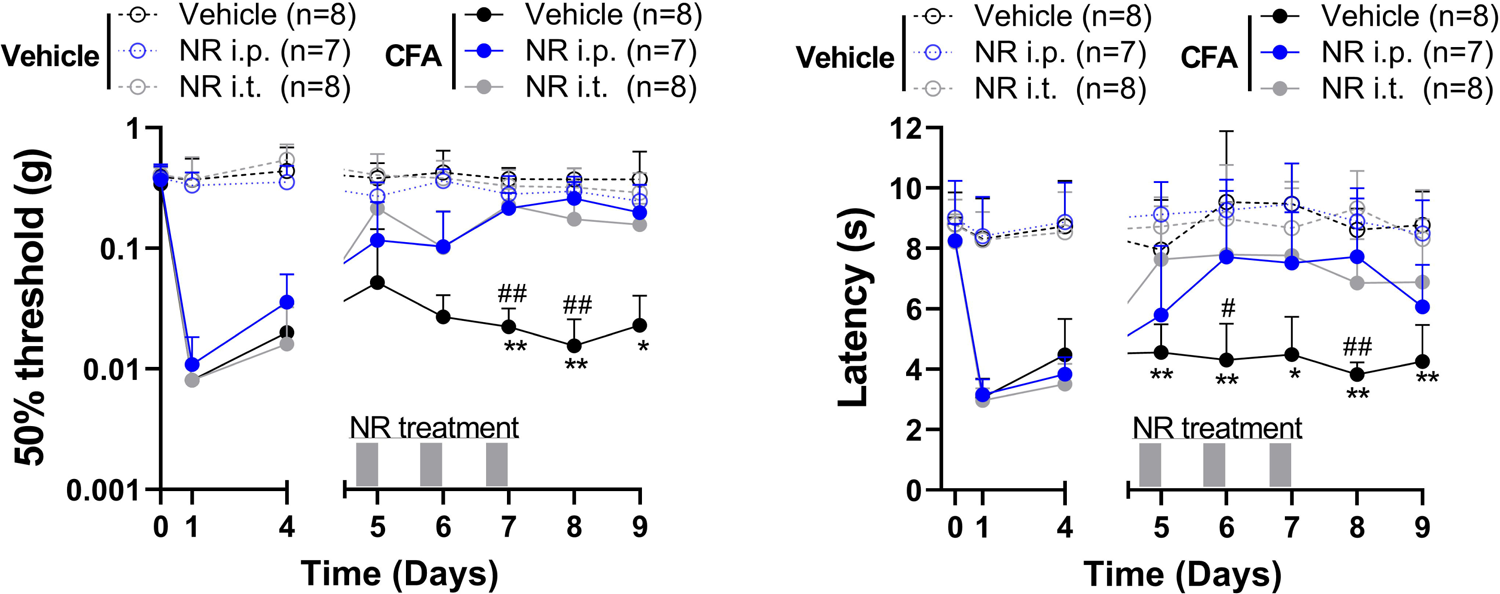
NAD^+^ supplementation attenuates chronic pain. **A/B)** Mice received intraperitoneal (i.p. 500 mg/kg) or intrathecal (i.t. 50 μg) injections with nicotinamide riboside (NR) at day 5, 6 and 7 after intraplantar complete freund’s adjuvant (CFA) or vehicle (* i.t. vehicle vs NR, # i.p. vehicle vs NR). **A)** Mechanical and **B)** thermal hyperalgesia was measured 4 hours after each NR administration. Data are represented as mean ± SD. *P < 0.05, **P < 0.01, ***P < 0.001. Statistical analyses were performed by two-way repeated measures ANOVA followed by a post-hoc Sidak’s multiple comparison test.

## Discussion

The mechanisms that impair resolution of inflammatory pain leading to persisting pain are still poorly understood. We identified that after resolution of inflammatory hyperalgesia, when latent plasticity of the sensory system is present, mitochondrial and metabolic changes persist in the DRG. These changes pointed to a disturbed redox balance. Importantly, these mitochondrial and metabolic disturbances are fundamental to the inability of primed mice to resolve from PGE_2_-induced inflammatory hyperalgesia (Figure 6). Notably, mitochondrial hyperactivity, metabolic disturbances, and priming-induced failure in resolution of PGE_2_-induced hyperalgesia were fully mimicked by increasing the expression of *ATPSCKMT* in DRG neurons. Attenuating mitochondrial hyperactivity, scavenging mtROS or NAD^+^ supplementation were sufficient to restore failed resolution of PGE_2_-induced hyperalgesia, and even promote pain resolution in a model of persistent inflammatory pain. Overall, these results highlight the importance of tight control of mitochondrial and metabolic activity in DRG neurons to ensure resolution of inflammatory pain.

**Figure 6:**
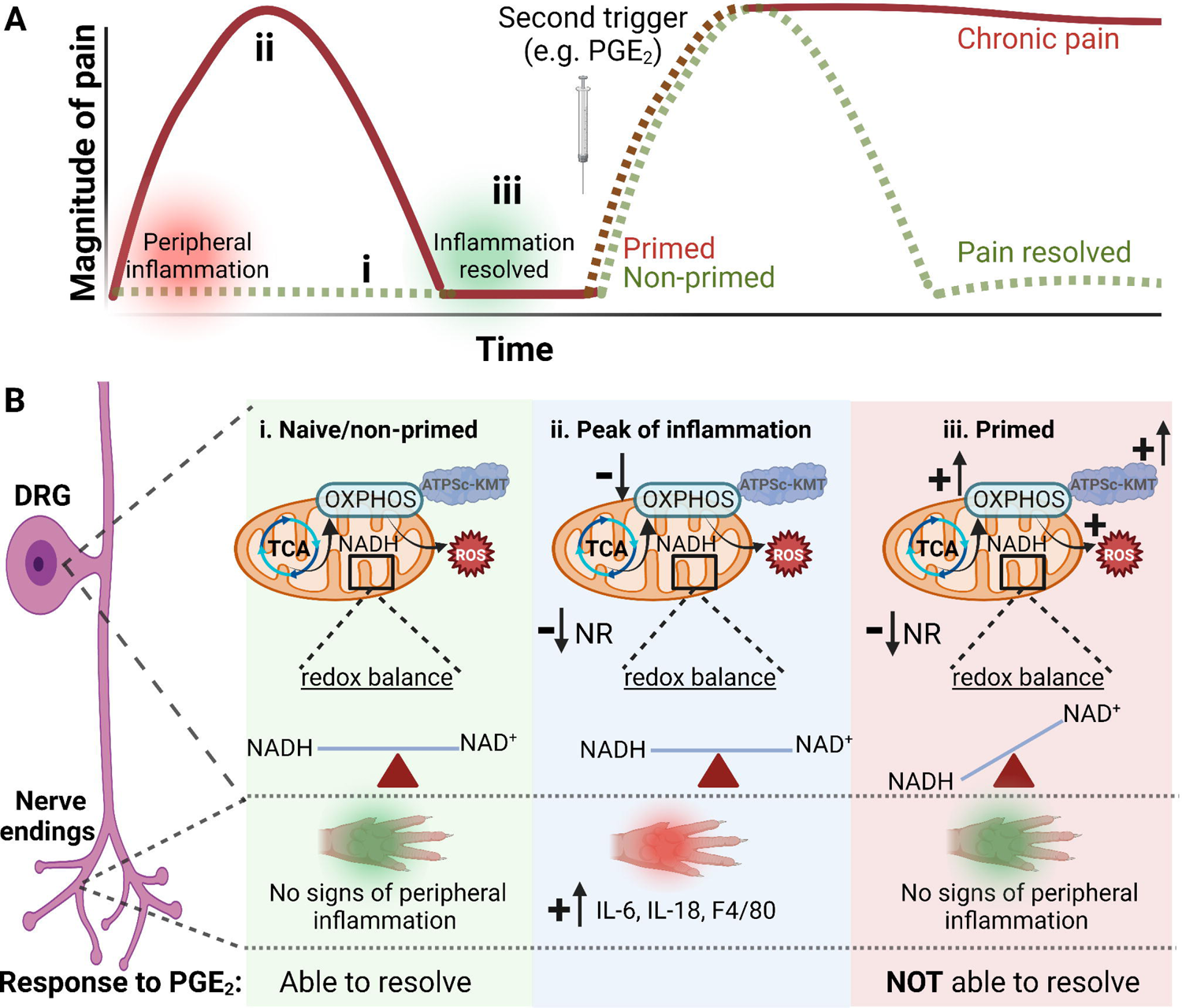
Schematic overview of proposed mechanisms. **A)** Time course of mechanical hypersensitivity in (i) naive mice and after intraplantar carrageenan, at the peak of (ii) acute pain and (iii) when hyperalgesia has resolved (primed state). A second stimulus (PGE_2_) is injected in primed and non-primed mice, where inflammatory hyperalgesia fails to resolve when mice were primed. **B)** A summary of our main findings at the level of DRG and paw at the different stages. Compared to naïve mice (i), at the peak of transient inflammatory pain (ii) there is a clear increase in inflammatory markers in the inflamed paw and a reduction in OXPHOS without disturbance in redox balance in the DRG. iii) When carrageenan-induced inflammatory hyperalgesia has resolved, signs of paw inflammation have disappeared. At this stage, ATPSc-KMT expression, and OXPHOS are increased in DRG neurons, concurrent with signs of oxidative stress (ROS production), reduced NR levels, and an increased mitochondrial NAD^+^/NADH ratio, possibly as a consequence of increased NADH consumption by enhanced activity of ETC. These peripheral-inflammation induced mitochondrial and metabolic disturbances in DRG lead to the inability to resolve from a subsequent inflammatory stimulus, like PGE_2_. Arrows with a plus (+) symbol indicate increase/upregulation, while the arrows with a minus (-) symbol suggest reduction in expression/activity. Figure created with BioRender.com.

Priming induced by a peripheral transient inflammation causes neuroplasticity of nociceptors, which includes increased mRNA translation and switch in cAMP signalling towards an PKCε-dependent pathway.^[9a, 30]^ The involvement of mitochondria in hyperalgesic priming has been previously postulated, because several mitochondrial proteins are targets of PKCε signalling.[31] However, earlier studies have excluded a potential role of mitochondria in peripheral nociceptor endings to restore failure in PGE_2_-induced hyperalgesia.[32] Nonetheless, mitochondria in the soma of nociceptors during hyperalgesic priming were not investigated. We hypothesized that the excitable soma of sensory neurons[33], is the ideal place to integrate signals and process axonal activity to initiate (transcriptional) adaptations to support altered neuronal functioning. Given this central role of the soma and the fact that mitochondria are mainly formed at the soma[34], mitochondrial fitness in the soma is key to maintain bioenergetics and cellular homeostasis and ensure the ability to adequately respond to noxious triggers. Indeed, we identified that a peripheral inflammation reduced mitochondrial respiration in both the soma of DRG neurons and sciatic nerves. However after resolution of hyperalgesia and inflammation, mitochondrial respiration was increased in the soma, but not in the axons. Moreover, inhibition of mitochondrial activity in the DRG, but not in the hind paw with nerve endings, restored resolution of inflammatory pain. Thus these results support the hypothesis that mitochondria are distinctly regulated in the soma of DRG neurons compared to sciatic nerves and their peripheral endings and mitochondria in the soma play an unique role in hyperalgic priming.[35]

We identified that DRG neurons maintain a higher mitochondrial respiration after resolution of a peripheral inflammation. Why would neurons benefit to do so? Various cell types, such as stress-induced murine and human fibroblasts, DRG neurons and peripheral blood mononuclear cells of diabetes mellitus or rheumatoid arthritis patients with an active disease increase their OXPHOS, possibly to compensate for a transient decline in NAD^+^ or ATP, to prevent oxidative stress and to promote cellular homeostasis.[36] The observed persistent increase of OXPHOS in DRG neurons of primed mice, possibly promotes the generation of ATP to regulate energy consuming processes, such as transport of proteins/organelles along cytoskeleton or to support increased protein translation, processes all needed to maintain the primed state.[37] Therefor we hypothesize that the inflammation-induced persisting increase in sensory neuron OXPHOS is likely needed to maintain the newly introduced homeostatic set-point of ‘primed’ neurons’ to permit this kind of ‘neuronal memory’ to respond differently to a future stimulus.

The precise mechanisms that maintain the enhanced OXPHOS in sensory neurons after priming is not fully understood. However, we found that carrageenan induced an increase in ATPsc-KMT expression in sensory neurons, which may be sufficient to increase OXPHOS in sensory neurons.^[8b]^ Similarly, ATPSc-KMT mRNA expression in DRG is also increased in a model of CFA-induced persistent inflammatory pain that is attenuated by either inhibition of complex I of ETC or *ATPSCKMT* knockdown.^[7b, 8a]^ These findings suggest that upregulating ATPSc-KMT expression occurs more broadly and is linked to changes in mitochondrial respiration, possibly to support cellular energy requirements after an inflammatory trigger.

The question arises how ATPsc-KMT promotes OXPHOS activity. ATPsc-KMT methylates Lys-43 of ATP synthase c-subunit (complex V of ETC) thereby regulating OXPHOS activity in several cell lines and primary neurons.^[8b]^ However, Lys-43 was already fully methylated in DRG of naïve mice, making it unlikely that ATPsc-KMT enhanced the OXPHOS via Lys-43 methylation in DRG neurons of primed mice. Possibly, ATPsc-KMT enhances OXPHOS by interacting and methylating other substrates. Indeed, when ATP synthase was fully inhibited and uncoupled from OXPHOS, *ATPSCKMT* deficient (ATPSc-KMT^-/-^) HAP1 cells still had a significantly reduced OCR compared to wild-type situation. This suggest that possible other ATPSc-KMT substrates are involved in regulating OXPHOS, something that remained to be elucidated.

Alterations in OXPHOS often generate ROS as a byproduct which may result in oxidative stress and a disturbed redox balance.^[36a, 38]^ We show here for the first time that disturbed mitochondrial NAD^+^/NADH redox balance persists in the DRG beyond resolution of inflammation. But how are changes in the redox state associated with pain? It is well known that a drop in cellular NAD^+^ level has detrimental effects for many biological processes. For example, a decline in NAD^+^ is associated with several pathologies, including neurodegenerative diseases.[15] In models of nerve injury and chemotherapy-induced chronic pain, NAD^+^ levels are reduced in sciatic nerves and DRG neurons.[39] In DRG lysates of primed mice, the NAD^+^ precursor nicotinamide riboside (NR) was reduced and the mitochondrial redox balance was disturbed. These changes occurred in parallel with increased mtROS levels in sensory neurons of primed mice. Thus possibly ROS production drives the excitability of sensory neurons. Indeed, ROS induce long term potentiation in neurons and/or sensitize neurons through transient receptor potential (TRP) channels, e.g. TRPA1 which is involved in mechanical and thermal (cold) hypersensitivity.[40] Another possibility is that changes in cellular NAD^+^ concentrations directly affect neuronal excitability because reduced cellular NAD^+^ levels inactivates potassium channels (e.g. the Slack K_NA_ channel) expressed in DRG neurons, which is associated with enhanced responsiveness of nociceptors to noxious stimuli.[41]

A limitation of this study is that we measured the metabolic profile in total DRG lysates, because it is technically difficult to measure metabolic profile specifically in isolated DRG neurons. Thus, we cannot fully exclude the possibility that observed metabolic differences occurred in cells other than DRG neurons, such as satellite cells or immune cells. Importantly, the numbers of immune cells in the DRG of primed mice (day 7 after carrageenan) did not differ from that in DRG of naïve mice^[3d]^, which supports that the changes in metabolites and redox balance are likely not caused by changes in the number of immune cells in the DRG, but rather by metabolic changes within DRG neurons. Although, we cannot fully exclude that metabolic alterations occurred within other cells in the DRG, sensory neurons likely contribute the most to these changes, also because they constitute the majority of cells in the DRG.[42] Moreover, ATPSc-KMT expression, glycolysis and mtROS formation were increased specifically in DRG neurons, suggesting that sensory neurons are the main contributors for the observed mitochondrial and metabolic changes within the DRG.

Various studies describe that NAD^+^ precursors, including NR, protect neurons when they are injured (e.g. Parkinson, Alzheimer’s disease).[43] However, whether NAD^+^ or its precursors have analgesic potential is less clear. Here we observed that supplementation of the NAD^+^ precursor NR was sufficient to restore pain resolution not only in primed mice, but also in CFA-induced persistent inflammatory pain, a model known to be associated with mitochondrial abnormalities in the DRG.^[7b]^ Oral administration of NR also reverses chemotherapy-induced and diabetic-induced neuropathy in rodents.^[39b, 44]^ There are several mechanisms that could explain how NAD^+^ promotes mitochondrial fitness and cellular homeostasis. The most described mechanism of action of NAD^+^ supplementation is the regulation of redox balance and antioxidant activity.^[14, 18a, 45]^ Another proposed mechanism is that NR improves mitochondrial fitness by activating cellular metabolic NAD^+^ sensors, such as sirtuins (NAD^+^ dependent histone deacetylase), which promote expression of antioxidant genes and genes involved in mitochondrial biogenesis.[46] Moreover, NR promotes mitophagy, i.e. removal of damaged mitochondria, in a neurodegenerative mouse model.^[19b]^ Thus, NR supplementation possibly facilitates mitochondrial quality control mechanisms in neurons that may overcome the observed priming-induced mitochondrial alterations. Overall, supplementation with NAD^+^ precursors, including NR, may have clinical benefit to treat chronic pain. Recent pharmacokinetic studies already proved that NR is safe and well-tolerated in humans.[45, 47] A small pilot study from 1959, described that NAD^+^ therapy induced remission and reduced pain-outcomes in arthritis patients.[48] However, a follow-up pilot study in 1996 described that nicotinamide treatment did not change pain levels, but improved joint flexibility, reduced inflammation which allowed osteoarthritis patients to reduce their anti-inflammatory medication.[49] Larger clinical trials are needed to investigate whether NAD^+^ supplementation may hold promise for treatment of chronic pain in inflammatory diseases.

In conclusion, here we provide evidence that a peripheral inflammation induces persistent mitochondrial and metabolic changes in the soma of sensory neurons, which affected the ability to resolve from hyperalgesia induced by a subsequent inflammatory trigger. Thus metabolic changes in sensory neurons result in failure of endogenous pain resolution pathways and drive the transition to chronic pain. Importantly, targeting mitochondrial respiration, scavenging ROS or supplementing NR represent a potential therapeutic strategy to restore failing pain resolution pathways to treat chronic inflammatory pain.

## Acknowledgements

We would like to thank Prof B. Burgering (UMC Utrecht) for access to the Seahorse instrument; R. van Eck and I. Aitink for their help with the analysis of immunofluorescence pictures; L. v. Vliet for complex-specific OCR measurements and Dr. F. Zwartkruis for help in designing CRISPR/CAS mediated ATPSC-KMT N2A cells. This work has received funding from Netherlands Organization for Scientific Research (H. Willemen; NWO, 016.VENI.192.053) and European Union’s Horizon 2020 research and innovation programme under the Marie Sklodowska-Curie grant agreement No 814244.

## Materials & Methods

### Animals

Experiments were conducted using adult male and female (aged 8-16 weeks) C57BL/6 mice (Janvier laboratories). Mice were maintained in the animal facility of the University of Utrecht and housed in groups under a 12h:12h light:dark cycle, with food and water available *ad libitum*. The cages contained environmental enrichment, including tissue papers and shelter. All experiments were performed in accordance with international guidelines and approved by the experimental animal committee of University Medical Center Utrecht (2014.I.08.059) or by the local experimental animal welfare body and the national Central Authority for Scientific Procedures on Animals (CCD, AVD115002015323 & AVD11500202010805).

Mice received an intraplantar injection (unilateral or in both hind paws) of 5 μl λ−carrageenan (primed, 1% w/v, Sigma-Aldrich) to induce transient inflammatory hyperalgesia. Non-primed mice received an intraplantar injection with saline. Day 7 after carrageenan or saline, mice were injected intraplantar with PGE_2_ (100 ng/paw, Sigma-Aldrich). For the induction of chronic inflammatory pain, mice received an unilateral intraplantar injection with 20 μl Complete Freund’s Adjuvant ((CFA), Sigma-Aldrich). Heat withdrawal latency times were determined using the Hargreaves test (IITC Life Science).[50] Mechanical thresholds were determined using the von Frey test (Stoelting) with the up-and-down method previously described.[51] In experiments in which mice received only intraplantar injections, each paw was considered an independent measurement in terms of latency times and 50% thresholds. In experiments with intrathecal or intraperitoneal drug administration, each animal was considered as independent measurement, so the average of the left and right paw were used. To minimize bias, animals were randomly assigned to the different groups prior to the start of experiment, and all experiments were performed by operators blinded to the treatments and/or genotypes.

### Drug administration

NAD^+^ precursor, nicotinamide riboside (NR) was injected intraperitoneal (500 mg/kg, Tebu-bio)[52] or intrathecal. In the CFA experiment, we measured hyperalgesia 4 hours after injection of NR. Intrathecal injections (5 μl) with NR (50 μg), myxothiazol (50 uM, Sigma-Aldrich), mito-tempol[23] (25 μg, Sanbio) and oligodeoxynucleotide (3 μg/μl day 4, 5 and 6 after carrageenan, Sigma-Aldrich), were performed under light isoflurane anesthesia as described.^[8a, 53]^ The following phosphorothioated sequences were used to specifically target mouse ATPsc-KMT: *ATPSCKMT*: CCCgCCTgTCTTTCTTCCTC *MM*: CgCCTCCgTTCCTTTCTCCT

### DNA and viral constructs

ATPSCKMT (NM_199133.4) was cloned in pIRES2-AcGFP1 (Clontech). We generated a bicistronic herpes simplex virus (HSV) construct expressing ATPsc-KMT and GFP as described previously.^[8a]^ Control empty HSV (HSV-EV) only expressed GFP. HSV was produced as previously described.[54] Mice were inoculated twice (day −3 and day −1 prior to PGE_2_) intraplantar with 2.5 μl of 1.4 × 10^7^ pfu/ml.

Lentivirus expressing shRNA against mouse *ATPSCKMT* was produced according protocol (Sigma-Aldrich). In short, 2×10^6^ HEK293T cells were cultured in Ø10 cm petri dish and transfected with PEI and a mix of plasmids, containing 5 µg MISSION® shRNA *ATPSCKMT* (or scrambled control), 1.8 μg envelope vector (pMD2.G) and 1.8 μg packaging vector (psPAX2). The medium was replaced the following day. Subsequently, 48 and 72 hours after transfection, medium containing lentivirus was collected for experiments.

### Cell lines, primary cell cultures and transfections

Mouse neuroblastoma Neuro2a (N2A) cells were kept in Dulbecco’s Modified Eagle medium (DMEM) with Glutamax-l containing 4,5g/L D-Glucose, pyruvate, 1% Penicillin/Streptomycin and 10% fetal calf serum. HAP1 cells were cultured in Iscove’s Modified Dulbecco’s Medium (IMDM) containing 10% fetal calf serum and 1% Penicillin/Streptomycin (P/S). *ATPSCKMT* overexpression was achieved with the plasmid described above, using Lipofectamin 2000 (Life technology) according to manufacturer’s instructions.

DRG were collected and primary sensory neurons were cultured as described.[55] Twenty-four hours after plating, sensory neuron cultures were inoculated with HSV (MOI of 2, 10,000 pfu) for 3 days. The anti-mitotic fluoro-deoxyuridine (FDU 13.3 μg/ml, Sigma-Aldrich) was added to inhibit satellite glia cell growth in the neuronal cultures. For *ATPSCKMT* knockdown, primary sensory neurons were incubated with lentivirus (50% of the medium) for 2 days.

### Mitochondrial superoxide detection with immunofluorescence

*In vivo*, MitoTrackerRedCM-H2XROS, which emits fluorescence upon oxidation, to measure mitochondrial superoxide production (10 μl of 100 μM, Life technology)^[8a, 21]^ was injected intrathecal at day 7 after intraplantar carrageenan administration. Six hours later, mice were perfused with PBS and 4% PFA and DRG were collected. Tissues were cryoprotected in sucrose, embedded in OCT compound (Sakura), and frozen at –80°C. Cryosections (10 μm) of lumbar DRG were stained with Neurotrace 435/455 (1:300, ThermoFisher) to visualize the neurons. For mitochondrial superoxide production measurements *in vitro*, HAP1 cells were incubated with 200 nM MitoTrackerRedCM-H2XROS in HBSS for 30 minutes. After PBS washes cells were fixed with 4% PFA. Fluorescence was captured using Olympus IX83 microscope (Olympus) microscope and analyzed with ImageJ software. The number of positive MitoTrackerRedCM-H2XROS neurons was determined by the amount of cells that had a higher fluorescence than the mean + 2x SD of the naïve group.

### Flow cytometry analysis for measuring mitochondrial mass

DRG (L3–L5) were collected from non-primed and primed mice. Tissues were gently minced and digested at 37 °C for 25[minutes with an enzyme cocktail (5 mg collagenase type I with 2.5 mg trypsin, Sigma-Aldrich) in 5 ml DMEM. Cells were incubated with 100 nM nonyl acridine orange (NAO, Enzo LifeSciences) for 30 min at 37°C to detect mitochondrial mass. After washing with PBS, the cells were incubated with ef506 viability marker (1:1000, eBioscience) for 20 min at 4°C, followed by a CD45-APC-ef780 staining (1:600, eBioscience) for 20 min at 4°C, to exclude the immune cells. Samples were acquired by Canto II flow cytometer (BD Biosciences) and analysed with FACSDIVA software.

### Mitochondrial bioenergetics assessment

N2A cells were plated (12.500 cells/well) on poly-l-lysine-coated XF24 wells plates (Seahorse Bioscience) in DMEM (high glucose, with pyruvate) (Gibco) + 10% FCS + 1% P/S. Primary DRG neurons (15K) were seeded on poly-d-ornithine/laminin coated XF24 wells plate, grown overnight at 37°C and transduced with HSV or lentivirus (see above). The cells were washed and placed in Seahorse XF-assay media (pH 7.4) containing 25 mM glucose, 4 mM glutamine and 1 mM pyruvate at 37°C for 1 h. The Seahorse Bioscience XFe24 Analyzer (Seahorse Bioscience) was used to measure oxygen consumption rate (OCR). After assessing basal OCR, 2 μM oligomycin, 2 μM FCCP and 2 μM of Rotenone and Antimycin A (all Sigma-Aldrich) were injected after cycle 3, 6, and 9 respectively. Each cycle consisted of 1.5 minute of mixing, 2 minutes waiting, and 3 minutes of measurements. For each condition three cycles were used. Glycolytic capacity was measured by subtracting the extracellular acidification rates after addition of oligomycin from basal ECAR levels. The measurements were normalized for protein content.

Mitochondria from N2A cells expressing ATPsC-KMT or control EV were isolated according to Iuso et al.[56] To measure complex II driven respiration, 5 μg of mitochondria were added in a non-coated XF24 plates in MAS buffer (220 mM d-Mannitol, 70 mM sucrose, 10 mM KH2PO4, 5 mM MgCl2, 2 mM HEPES, 1 mM EGTA, and 0.2% (w/v) of fatty acid-free BSA, pH 7.2) supplemented with 10 mM succinate and 2 µM rotenone. OCR levels were measured under basal conditions, and after sequential addition of ADP (2 mM), oligomycin (3,2 μM), FCCP (4 μM), and antimycin A (4 μM), to measure state III. Each assay cycle consisted of 1 minute of mixing and 3 minutes of OCR measurements. For each condition, three cycles were used to determine the average OCR under given condition.

Measurement of mitochondrial respiration in sciatic nerves was performed according to Kruwoski et al.[57] In short, sciatic nerves were isolated and stored for maximum 1 hr on ice in XF media, until all nerves were collected. Sciatic nerves were transferred into islet capture XF24 microplates containing XF-assay media supplemented with 5.5 mM glucose, 0.5 mM sodium pyruvate, and 1 mM glutamine (pH 7.4). Plates were incubated in a non-CO_2_ incubator to degas for 2 hrs at 37°C. OCR levels were measured under basal conditions and after sequential addition of oligomycin (12 μM), FCCP (20 μM), and antimycin A/Rotenone (20 μM). Each assay cycle consisted of 3 minute of mixing, 3 minutes waiting, and 4 minutes of measurements. For each condition four cycles were used. The measured OCR was normalized for protein content.

N2A cells were used to measure complex-specific respiration. To assess complex I activity, cells were incubated with pyruvate and malate. To assess complex II/III activity, cells were incubated with succinate and rotenone to supply electrons to the ETC via complex II and rotenone to inhibit complex I. To assess complex IV activity, cells were incubated with TMPD, ascorbate to supply electrons to the ETC via complex IV and antimycin A to inhibit complex II/III. All reagent solutions were prepared in MAS buffer (660 mM mannitol, 210 mM sucrose, 30 mM KH2PO4, 15 mM MgCl2, 6 mM HEPES and 3 mM EGTA), in Ultrapure or tissue-culture-grade H2O and adjusted to pH 7.2 at 37°C, unless stated otherwise. Ascorbate (1,33 M, Genfarma, Toledo, Spain), N,N,N′,N′-tetramethyl-p-phenylenediamine (10 mM; also known as TMPD, Sigma-Aldrich) dissolved in 10 mM ascorbate and ADP (50 mM, Sigma-Aldrich) were freshly prepared. All other reagents (Sigma-Aldrich) were prepared beforehand and stored at −20°C, including pyruvate (1 M), succinate (0.5 M), malate (0.5 M), dichloroacetic acid (1 M; DCA), rotenone (2.5 mM), antimycin A (2.5 mM), myxothiazol (10 mM) and carbonyl cyanide-p-trifluoromethoxyphenylhydrazone (2.5 mM; FCCP). The medium was removed and cells were washed once with MAS buffer. MAS buffer containing 10 mM pyruvate, 10 mM malate, 2 mM DCA, 4 mM ADP, 1 nM XF PMP (Seahorse Bioscience) and with 2.5 mM FCCP and 2.5 mM oligomycin was added to the cells. The cartridge was calibrated one day prior to the experiment following manufacturer’s instructions. The cartridge was loaded with 75 μl of the following solutions: (port A) Rotenone (2 μM) in MAS buffer, (port B) succinate (10 mM), rotenone (2 μM) in MAS buffer, (port C) antimycin A (2 μM) in MAS buffer and (port D) ascorbate (10 mM), TMPD (100 μM), antimycin A (2 μM) in MAS buffer. The OCR was measured using the following steps: mix/delay/measure times were 0.5 min/0.5 min/2 min, an equilibration step was not included and three measurements were made for each step. OCR was recorded as pM/minute.

### Immunostaining

DRG were collected and directly embedded in OCT compound (Sakura) and frozen at –80°C. Cryosections (10 μm) of lumbar DRG were post-fixed in PFA and stained with anti-ATPSc-KMT (1:500, biorbyt). ATPSc-KMT was visualized by using alexafluor 594-conjugated secondary antibody. Neurons were visualized with Neurotrace 435/455 (1:300, ThermoFisher). Photographs were captured using an Olympus IX83 microscope using identical exposure times for all slides within one experiment. Fluorescence intensity was analyzed with ImageJ software. Fluorescence was analyzed in small diameter neurons < 20 μm and medium/large diameter neurons > 20 μm.[22]

### Collection of cell lysates for metabolomics

HAP1 expressing wtATPSc-KMT (WT), ATPSc-KMT^-/-^ and reconstituted with WT and E117A mutated ATPSc-KMT were plated in 6-well plates and cultured for 48 hours. Medium was refreshed 24 hours after plating. Cell collection was done by washing cells with cold PBS (4 °C), followed by cell scraping in 0.5 ml ice-cold methanol. Next, methanol samples were transferred into Eppendorf tubes, centrifuged (13000 rpm for 5 min at 4 °C), and then supernatants were transferred to new 1.5-ml Eppendorf tubes. The samples were evaporated at 40 °C under a gentle stream of nitrogen until complete dryness, and reconstituted with 500 μl of UPLC-grade methanol (room temperature). The reconstituted samples were stored at − 80°C until analysis was performed.

### Non-quantitative direct-infusion high-resolution mass spectrometry (DI-HRMS)

A non-quantitative DI-HRMS metabolomics method was used as previously described (Haijes et al. 2019). Samples were analyzed using a TriVersa NanoMate system (Advion, Ithaca, NY, USA) controlled by Chipsoft software (version 8.3.3, Advion). Data acquisition was performed using Xcalibur software (version 3.0, Thermo Scientific, Waltham,MA, USA). DI-HRMS is unable to separate isomers, therefore mass peak intensities consisted of summed intensities of these isomers. Metabolite annotation was performed using a peak calling pipeline developed in R programming language, annotated the raw mass spectrometry data according to the Human Metabolome DataBase (HMDB) allowing for 2 ppm deviation from the theoretical m/z. This resulted in ∼3800 metabolite annotations corresponding to ∼1900 unique metabolite features. Data was analyzed with MetaboAnalyst 5.0. In case of multiple possible annotations per feature, isobaric compounds were processed as only one metabolite for statistical purposes.

### Ultra-high performance liquid chromatography (UPLC)

Liquid chromatography coupled to tandem mass spectrometry (LC/MS-MS) was performed using a Q Exactive™ HF hybrid quadrupole-Orbitrap mass spectrometer (Thermo Fisher Scientific). In short, cells from a single well of a 6-wells plate were quenched using 250μL of −80°C 80:20 methanol/water. Next, cells were incubated on dry ice for 20 minutes and scraped on ice, subsequently, the extract was collected in 1.5mL Eppendorf tubes. The Eppendorf’s containing the extract were then vortexed and centrifuged at 14000 G, after that the supernatant was collected in a new Eppendorf tube. Extracts were dried under a flow of nitrogen at 40°C, reconstituted in 40μL of water after which 10μL of 100μM β-NAD-d4 (Toronto Research Chemicals) was added resulting in a final concentration of 20μM for the internal standard (IS). Prior to analysis, calibration samples were prepared by dilution and subsequent addition of the internal standard. To this end, standards were serially diluted in a range of 125μM to 3.9μM (NAD+) and 15.5μM to 1μM (NADH). Sample extracts and standards were transferred to 12×32mm glass screw neck vials (Waters) and then injected onto a Sunshell RP-Aqua column(150 mm × 3 mm i.d., 2.6 μm; ChromaNik Technologies Inc., Osaka, Japan). For this purpose, a binary solvent gradient comprising of 0.1% formic acid in water (Mobile phase A) and 0.1% formic acid in acetonitrile (Mobile phase B) was used. The flow rate was 0.6ml/min with the following gradient elution: isocratic 100% A from 0-0.5 min, linear from 100 to 85% A from 0.5-2 min, linear from 85 to 75% A from 2-2.75 min, linear from 75 to 30% A from 2.75-3.5 min, isocratic 30% A from 3.5-7 min, linear from 30 to 100% A from 7-7.2 min and isocratic 100 % A (initial solvent conditions) from 7.2 to 12 to equilibrate the column. The column flow was then directed into the MS detector and the samples were measured. Raw data integration and inspection were performed using TraceFinder 4.1 software (Thermo Fisher Scientific).

### Quantification TCA cycle intermediates

The quantification of TCA cycle intermediates in cell lysates was performed on a Waters Acquity ultra performance liquid chromatography system (Waters Corp., Milford, USA), as described by Broeks et al.[58] The assay was performed for technical triplicates of cell lysates and data were corrected for total protein concentration.

### Western Blot

Protein concentrations of the total cell lysates of lumbar DRG were determined using a Bradford assay (Bio-Rad). Protein samples (40 μg) were separated by 4-10% SDS-PAGE and transferred to a PVDF membrane (Immobilon-P, Millipore). Membrane was stained with anti-OXPHOS (1:1000, ThermoFisher scientific) or goat anti-β-actin (1:1000, abcam), followed by incubation with 1:5000 donkey anti goat-HRP (abcam). Specific bands were visualized by chemiluminescence (ECL, Advansta) and imaging system Biorad.

### Real-time (RT)-PCR

Total RNA from freshly isolated DRG or hind paws was isolated using TRizol and RNeasy mini kit (Qiagen). cDNA was synthesized using iScript reverse transcription supermix, according to manufacture protocol (Bio-Rad, Hercules, CA). Quantitative real-time PCR reaction was performed with a QuantStudio 3 (ThermoFisher) following manufacturer’s instructions. We used the following primers:

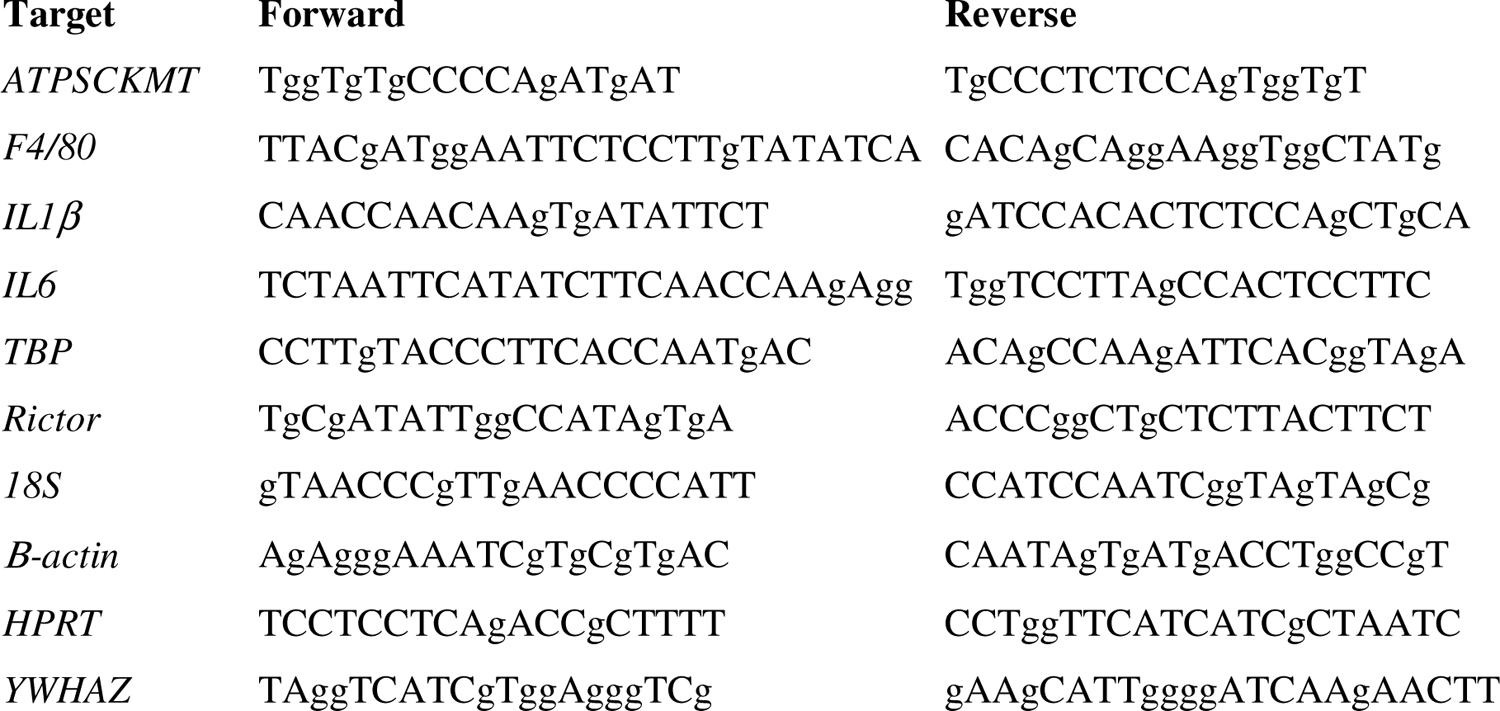

The most stable housekeeping genes per tissue were determined by measuring multiple internal controls (*β-actin, 18S, GAPDH, B2M, HRPT, β-tubulin, Rictor, TBP, YWHAZ*) and were analyzed with geNorm.[59] mRNA expression is represented as relative expression = 2^Ct(average^ ^of^ ^reference^ ^genes)-Ct(target).^ For mRNA expression in the paws, we used the average Ct values of *HPRT*, *β-actin* and *YWHAZ* as reference; for mRNA expression in DRG we used the average of *18S*, *TBP* and *Rictor* as reference.

### Preparation of DRG extract enriched for mitochondrial inner membrane proteins

Preparation of DRG extract enriched for mitochondrial inner membrane proteins was performed at 4 °C, similarly as described previously.^[8b]^ DRG, frozen in PBS, were thawed on ice, centrifuged (11000 x g, 1 min), and the PBS was discarded. The pellet was resuspended in 50 μl PBS, supplemented with 0.5 mg/mL digitonin and 1% protease inhibitor cocktail (P8340), and incubated for 5 min on ice. The suspension was centrifuged (11000 x g, 10 min) and the supernatant discarded. The pellet was resuspended in 50 μl of Extraction Buffer (50 mM Tris-HCl pH 7.4, 100 mM NaCl, 1% *n*-dedecyl-β-D-maltoside, 5% glycerol, and protease inhibitors), and incubated on ice for 5 min. The suspension was centrifuged (16100 x g, 5 min) and *n*-dedecyl-β-D-maltoside-extracted proteins were recovered in the supernatant.

### Mass spectrometry analysis of ATPSc from DRG

Proteins in DRG extracts were resolved by SDS-PAGE, stained with Coomassie, and the region of gel containing ATPSc, i.e. around the 8 kDa marker, was excised and subjected to in-gel chymotrypsin (Roche) digestion. The resulting proteolytic fragments were analyzed by liquid chromatography MS, similarly as described previously.[60] MS data were analyzed using an in-house maintained human protein sequence databases using SEQUEST^TM^ and Proteome Discoverer^TM^ (Thermo Fisher Scientific). The mass tolerances of a fragment ion and a parent ion were set as 0.5 Da and 10 ppm, respectively. Methionine oxidation, cysteine carbamido-methylation, lysine mono-, di- and trimethylation, and arginine mono- and dimethylation were selected as variable modifications. MS/MS spectra of peptides corresponding to methylated Lys-43 in ATPSc were manually searched by Qual Browser (v2.0.7).

### Statistical analysis

All data are presented as mean ± SD and were analyzed with GraphPad Prism version 9.3 using Student’s t-test (two-tailed, unpaired), Fisher-exact test, one-way, two-way ANOVA, or two-way ANOVA with repeated measures as appropriate, followed by post-hoc analysis. The used post-hoc analyses are indicated in each figure. A p-value less than 0.05 was considered statistically significant and each significance is indicated with *p<0.05, **p<0.01, ***p<0.001. The n is depicted in the figures, or in the figure legends.

## Supplementary information

Figures S1-S5

