## Supplementary figures and images for "Inflammation-induced mitochondrial and metabolic disturbances in sensory neurons control the switch from acute to chronic pain"

### Suppl Figure 1

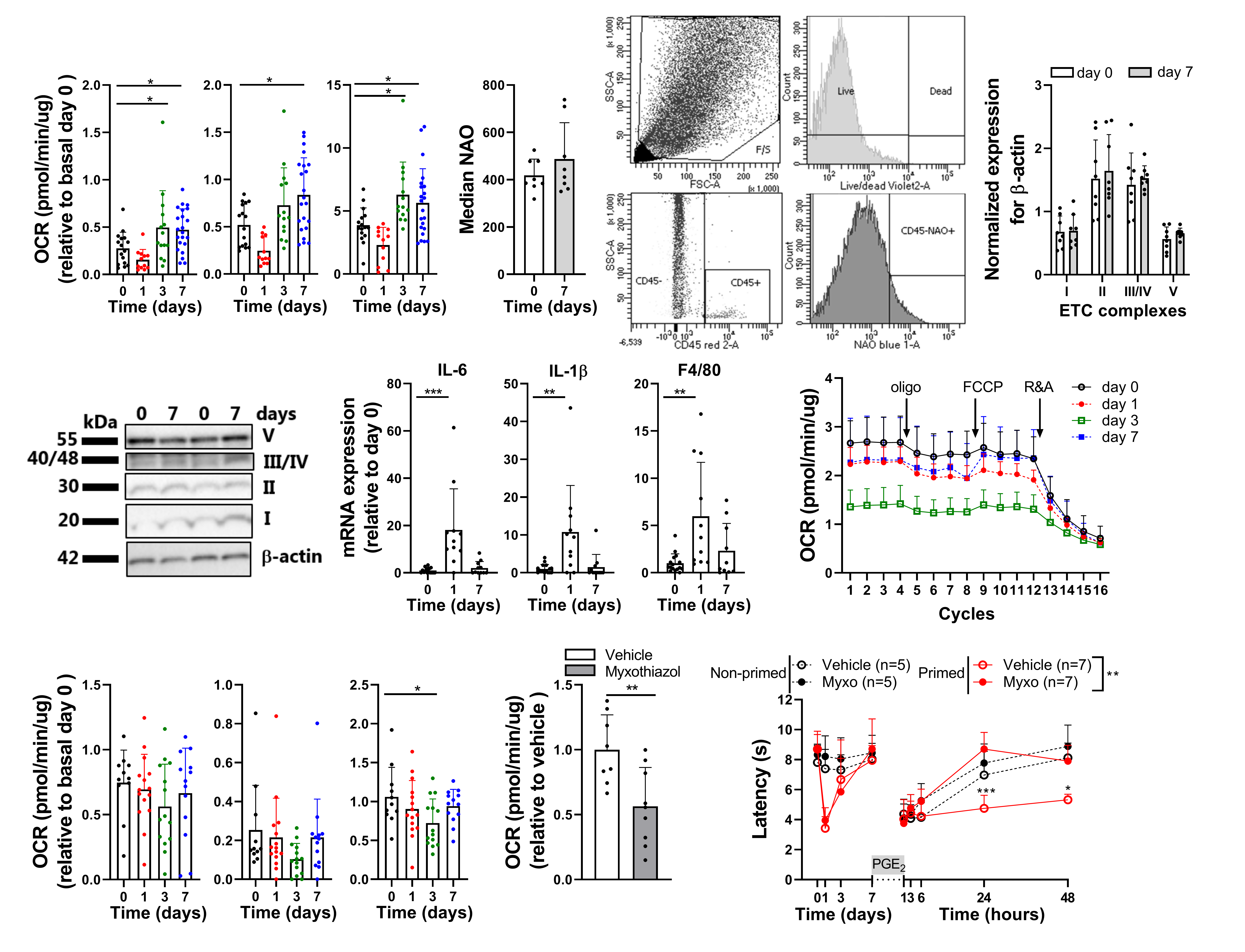

### Suppl Figure 2

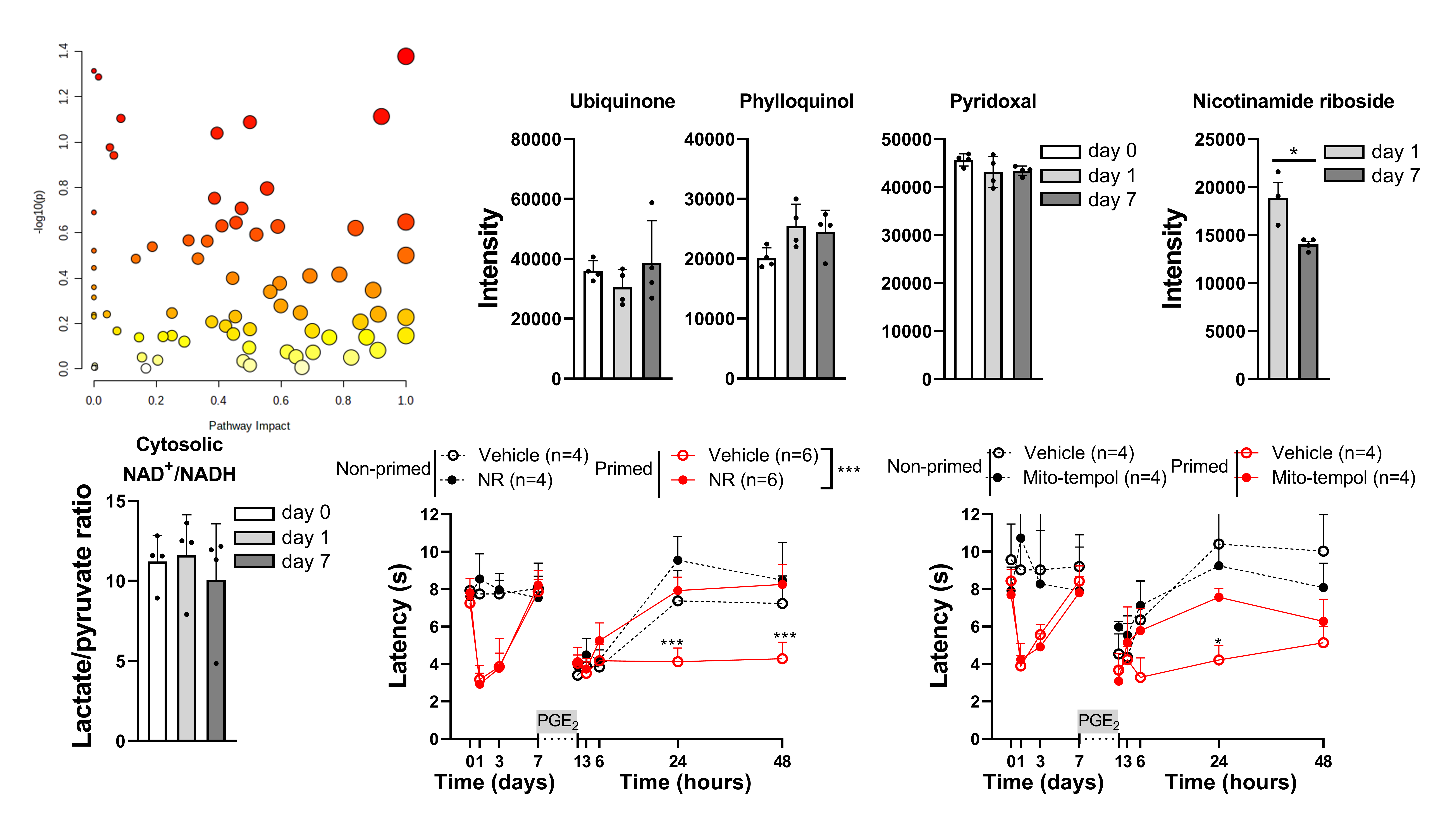

### Suppl Figure 3

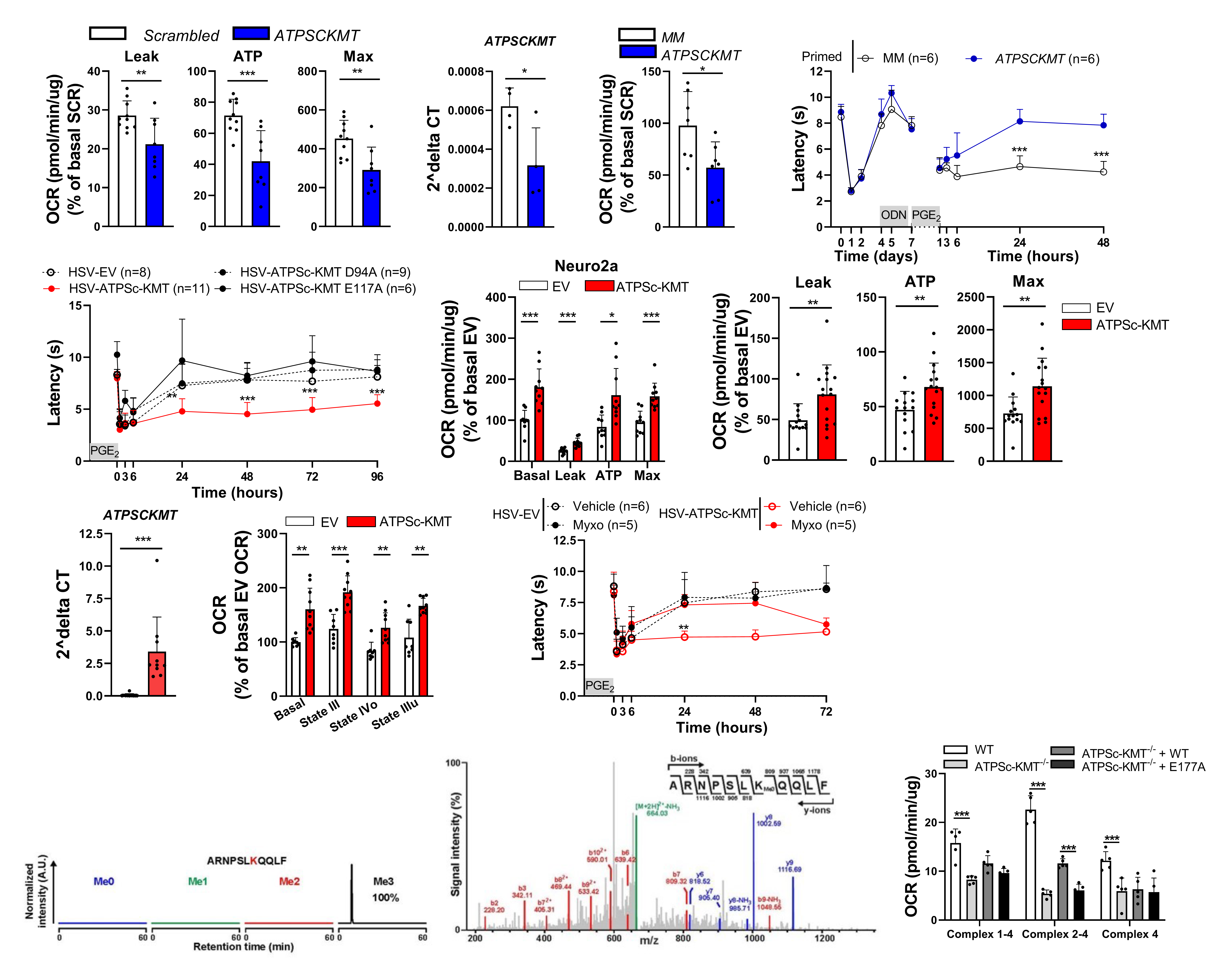

### Suppl Figure 4

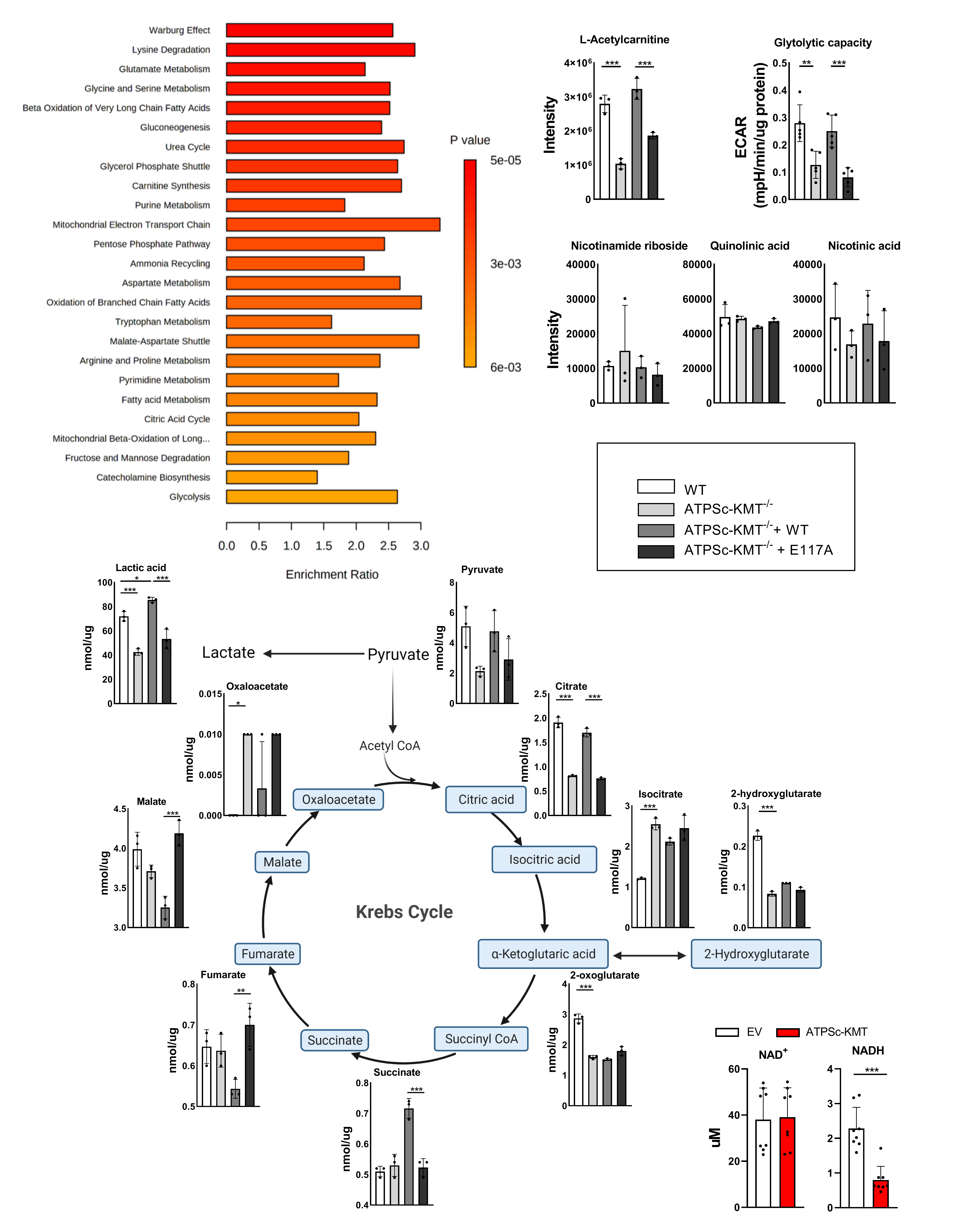

### Suppl Figure 5

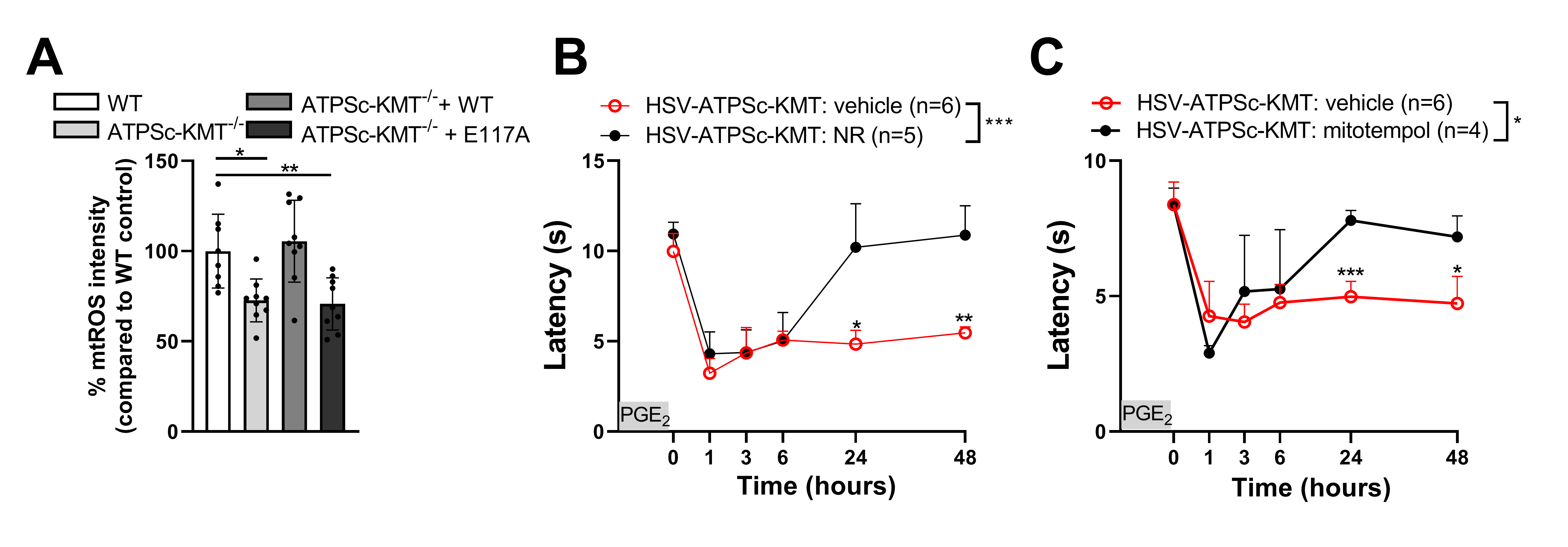
