## Supplementary material for "Inflammation-induced mitochondrial and metabolic disturbances in sensory neurons control the switch from acute to chronic pain": Suppl legends

**Supplementary figures:**

**Figure S1:** **A)** OCR in DRG neuron cultures at day 0 (n=16), 1 (n=12), 3 (n=14) and 7 (n=22) after intraplantar carrageenan injection. Leak, ATP-driven and maximal respiration was measured after sequential addition of oligomycin (ATP synthase inhibitor), carbonyl cyanide-p-trifluoromethoxyphenylhydrazone (FCCP, uncoupling protonophore that dissipates mitochondrial membrane potential), and a mixture of rotenone (inhibitor of Complex I) and antimycin A (inhibitor of Complex III). **B)** FACS quantification of nonyl acridine orange (NAO) positive CD45- cells in DRG neurons, as a measure of mitochondrial content in non-primed (day 0) and primed (day 7) mice (n=8). **C)** FACS gating strategy for measuring mitochondrial mass in B. **D)** Quantified expression of the five different OXPHOS complexes (normalized for *β-actin*) in DRG of primed (day 7) and non-primed mice (day 0) mice (n=8). **E** ) Exemplar image of the western blot that was used to quantify expression for D. **F)** Relative mRNA expression of inflammatory markers in the paw at indicated days after intraplantar carrageenan injection (n=6). **G**) OCR was measured overtime in sciatic nerves at day 0 (n=11), 1 (n=15), 3 (n=14) and 7 (n=13) after intraplantar carrageenan injection. **H)** Leak, ATP-driven and maximal respiration in sciatic nerves was measured, similar as in A. **I)** Basal OCR in DRG neuron cultures of primed mice (day 7) 1 hour after intrathecal injection of vehicle or myxothiazol (myxo, 50 μM) (n=8). **J**) Course of PGE_2_-induced thermal hyperalgesia after intrathecal injection of vehicle or myxothiazol (myxo, 50 μM) at day 7 in carrageenan-primed and non-primed mice, and 15 min prior to intraplantar PGE_2_. Data are represented as mean ± SD. *P < 0.05, **P < 0.01, ***P < 0.001. Statistical analyses were performed by, student’s T-test (B and I), one-way ANOVA (A, F, H) followed by Dunnett’s multiple comparison test or two-way ANOVA followed by a post-hoc Sidak’s multiple comparison test (D) and with repeated measures (J: stars indicate significance comparing primed conditions).

**Figure S2:** **A)** Pathway analysis of metabolites detected in lumbar DRG of untreated mice (day 0) versus DRG from primed mice (day 7 after carrageenan). Colors varying from yellow to red mean different levels of significance, in red are the most significant affected pathways indicated. Figure created with MetaboAnalyst 5.0. **B/C)** Intensity of metabolites in lumbar DRG **B)** involved in ubiquinone synthesis and vitamin B6 metabolism, and **C)** including nicotinamide riboside at indicated days (n=4). **D)** Lactate/pyruvate ratio as an indirect measure of cytosolic NAD^+^/NADH ratio (n=4). **E/F)** Course of PGE_2_-induced thermal hyperalgesia after **E)** intraperitoneal injection with nicotinamide riboside (NR, 500 mg/kg) or **F)** intrathecal administration of mito-tempol (25 ug) at day 7 in carrageenan-primed and non-primed mice and 15 min prior to intraplantar PGE_2_. Data are represented as mean ± SD. *P < 0.05, ***P < 0.001. Statistical analyses were performed by, student’s T-test (C), one-way ANOVA (B and D) followed by Dunnett’s multiple comparison test or two-way repeated measures ANOVA followed by a post-hoc Sidak’s multiple comparison test (E and F: stars indicate significance comparing primed conditions).

**Figure S3:** **A)** OCR measurements in DRG neuron cultures after lentiviral-mediated *ATPSCKMT* knockdown (n=8) compared to scrambled-controls (n=10). Leak, ATP-driven and maximal respiration was measured after sequential addition of oligomycin, FCCP, and mixture of rotenone and antimycin A. **B)** *ATPSCKMT* mRNA expression in DRG neurons after lentiviral-mediated knockdown compared to scrambled-controls (n=4). **C)** Basal OCR measurements in DRG neuron cultures at day 7 and after intrathecal injection of ATPSCKMT antisense or mismatched control (MM) at day, 4, 5 and 6 (n=7). **D)** Course of PGE_2_-induced thermal hyperalgesia after intrathecal ATPSCKMT antisense or MM oligodeoxynucleotide (ODN, 3 μg/μl, 5 μL) at day 4, 5 and 6 in carrageenan-primed mice. **E)** Course of PGE_2_-induced thermal hyperalgesia in mice expressing ATPSc-KMT, its enzyme-dead mutant (D94A or E117A), or control empty vector (EV) in DRG neurons. Intraplantar HSV injections were administrated at day -3 and -1 (35.000 pfu/paw) **F/G)** OCR measurements in **F)** N2A’s after overexpression of ATPSc-KMT or EV control (n=10), and **G)** primary sensory neurons after HSV-mediated expression of ATPSc-KMT (n=16) or EV (n=14); measurements were performed as described in A. **H)** ATPSCKMT mRNA expression in DRG neurons after transduction with HSV (n=10). **I)** OCR measurements in isolated mitochondria from N2A cells overexpressing ATPSc-KMT (n=10) or EV (n=8). Complex-II driven basal, StateIII, State IVo and StateIIIu was measured after sequential addition of ADP, oligomycin, FCCP and antimycin A. **J)** Same as E, but after intrathecal injection of myxothiazol (myxo, 50 uM) 15 min prior to intraplantar PGE_2_. **K)** Methylation states of the peptide ARNPSL**K**QQLF containing lysine-43 (bold) of the ATP synthase in DRG under naïve conditions. **L)** MS/MS fragmentation spectra shows that lysine-43 in ARNPSLK(me3)QQLF is tri-methylated in DRG. **M)** OCR measurements in permeabilized HAP1 cells in addition of FCCP/Oligomycin (n=5). Cells were incubated with pyruvate and malate to supply electrons to the ETC via complex I, succinate and rotenone to supply electrons to the ETC via complex II and TMPD, ascorbate and antimycin A to supply electrons to the ETC via complex IV. Data are represented as mean ± SD. *P < 0.05, **P < 0.01, ***P < 0.001. Statistical analyses were performed by Student’s t-test (A-C and G/H), one-way ANOVA (F and I) followed by Dunnett’s multiple comparison test or two-way repeated measures ANOVA followed by a post-hoc Sidak’s multiple comparison test (D/E, J and M: stars indicate significance comparing ATPSc-KMT-primed conditions). Primed mice by ATPSc-KMT overexpression are indicated with red bars/lines and blue bars/lines indicate *ATPSCKMT* knockdown.

**Figure S4**: **A/B)** Top 25 of enrichment analysis, visualizing **A)** mechanisms and **B)** an example of a functionally related metabolite that is significantly changed between ATPSc-KMT^-/-^ cells and WT cells (n=3). **C)** Targeted screen with focus on TCA metabolites, a-ketoglutaric acid also referred as 2-oxoglutarate (n=3). Figure created with BioRender.com **D)** ECAR, as measure for glycolytic capacity, in the 4 different cell conditions (n=5). **E)** Intensity of metabolites involved in generation of NAD^+^ (n=3). **F)** NAD^+^ and NADH pool in N2A’s after overexpression of ATPSc-KMT or EV control (n=8). Data are represented as mean ± SD. *P < 0.05, **P < 0.01, ***P < 0.001. Statistical analyses were performed by Student’s t-test (F) or one-way ANOVA (B-E) followed by Dunnett’s multiple comparison test.

**Figure S5**: **A)** mtROS formation in the four different cell conditions (n=9). **B/C)** Course of PGE_2_-induced thermal hyperalgesia after **B)** intraperitoneal injection with nicotinamide riboside (NR, 500 mg/kg) or **C)** intrathecal injection of mito-tempol (25 ug)) in mice expressing ATPSc-KMT in DRG neurons. Intraplantar HSV injections were administrated at day -3 and -1 (35.000 pfu/paw). Data are represented as mean ± SD. *P < 0.05, **P < 0.01, ***P < 0.001. Statistical analyses were performed by one-way ANOVA (A) followed by Dunnett’s multiple comparison test or two-way repeated measures ANOVA followed by a post-hoc Sidak’s multiple comparison test (B and C).
